# Noncoding and Coding Mechanisms of Aging Heart Failure with Preserved Ejection Fraction with Thyroid Dysfunction

**DOI:** 10.1101/2025.02.04.636476

**Authors:** Sankalpa Chakraborty, Bryce Dickerson, Curren Bounds, Sophia Lemus, Caleb Hickman, Viswanathan Rajagopalan

## Abstract

Heart Failure with preserved Ejection Fraction (HFpEF) is a heterogeneous geriatric syndrome with complex pathophysiology and comorbidities. Long noncoding RNAs (lncRNAs) account for a large majority of the functional mammalian transcriptome and act as key regulators in complex physiological and pathological processes. However, the role of lncRNAs and the development of thyroid dysfunction in aging HFpEF is not clear. We investigated the ZSF1 model in both early and more severe stages of HFpEF (5-, 13- and 20-months old [mo]). We assessed molecular, biochemical, and pathophysiological roles involving lncRNAs, mRNAs, and inflammatory markers. Thyroid hormone (TH) immuno-sorbent assays showed significant decreases in serum T3 levels in 5-mo and serum T4 levels in 5-mo and 13-mo obese HFpEF groups compared to lean controls. Morphometric analyses showed significant increases in heart and LV weights in obese HFpEF rats indicating cardiac hypertrophy. LncRNA microarray and RT-qPCR revealed that three key lncRNAs were significantly increased in 13-mo obese HFpEF but not in 5-mo obese HFpEF left ventricles compared to the ZSF1 lean controls. Microarray analyses showed that *Sik1* mRNA was significantly upregulated and *Anxa13 w*as downregulated in early obese HFpEF hearts compared to the lean controls. Additionally, we also uncovered previously unreported tissue and serum inflammatory cytokine profiles in early and late HFpEF. This study has identified key novel lncRNA and inflammatory markers in early and late hypothyroid HFpEF. Further studies may help in better understanding and development of diagnostic and therapeutic targets for HFpEF that presents with severe morbidity and mortality.

## Background

Heart Failure (HF) is considered a global pandemic affecting about 64 million people worldwide and about 6.7 million people in the US (1, 2). By the year 2030, HF is estimated to cost approximately $69.8 billion (1). The prevalence of HF is increasing, and this may be attributed to the increased aging of the population, increasing prevalence of HF risk factors and comorbidities, improved treatment/survival following cardiovascular (CV) diseases, increased awareness and improvement in diagnostic precision, etc. (1, 3).

Heart Failure with preserved Ejection Fraction (HFpEF) constitutes approximately 50% of HF cases (4, 5), with increasing prevalence over the past two decades (3, 6). HFpEF is characterized by HF with left ventricular (LV) ejection fraction (EF) greater than or equal to 50%. In many cases, it is also associated with abnormal LV diastolic function (7), comorbidities/risk factors including hypertension, metabolic and thyroid dysfunction, advanced age, inflammation, obesity, diabetes mellitus, and renal dysfunction (8–12). HFpEF remains a major public health concern with high morbidity and mortality rates in the United States and across (4), and treatment options are limited. Compared to HF with reduced EF or HFrEF, older age was more strongly associated with HFpEF relative risk per 10-year(13).

Thyroid hormones (THs) are crucial CV regulators. Both T3 (Triiodothyronine; active form) and T4 (Thyroxine; prohormone) are secreted by the thyroid gland. Hypothyroidism is generally associated with low serum TH levels, decreased contractility and heart rate (HR), impaired diastolic relaxation, slowing of metabolic processes, dyslipidemia, inflammation, arrhythmia, etc.(14–18). We and others have shown that THs may be decreased in serum and/or cardiac tissues in various forms of CV and associated disorders, and optimal T3 dosing improved cardiac performance in hypothyroid cardiomyopathy, diabetic cardiomyopathy, hypertension, HF, myocardial infarction, ischemia-reperfusion injury, etc. without major adverse effects (19–25).

With only about 2% of the genome translating to proteins, the vast majority of the transcribable transcriptome is comprised of noncoding RNAs (ncRNAs). The majority (∼93%) of disease- and trait-associated variants emerging from Genome-Wide Association Studies were reported to lie within the noncoding regions (26). Long non-coding RNAs (lncRNAs), which are defined as non-coding RNAs >200 nucleotides, comprise the most functionally diverse class of ncRNAs (27, 28). LncRNAs have been found to be critical for the development and progression of a variety of cardiovascular disorders. They have also been shown to be important in serving as biomarkers and therapeutic targets and a few are in clinical trials (29–31). However, the role of aforesaid noncoding RNAs in the development of hypothyroid (low thyroid hormonal [TH] levels), especially in the aging/aged HFpEF heart is not clear.

This study explores the mechanistic landscape of HFpEF from multiple standpoints – aging, lncRNAs, mRNAs, thyroid dysfunction, and inflammatory mediators. We used the Zucker fatty and Spontaneously Hypertensive (ZSF1) obese rat model that presents with relevant features of clinical HFpEF including but not limited to diastolic dysfunction, HF, insulin resistance, hyperglycemia, hypertension, hyperlipidemia, exercise intolerance, etc. While the obese are reported to have mutations in the leptin receptor (fa, cp), the lean counterparts can be heterozygous and may serve as useful controls presenting with hypertension, but without HF or obesity (11, 32). The findings presented in this study have not only shown transcripts associated with the early stage of HFpEF but also identified lncRNAs associated with aging and severity of the condition. In addition, this study also reports important pathways of coding RNAs along with multiple key markers of inflammation, which is a critical pathophysiological process in HFpEF.

## Methods

### Animal models

All protocols were approved by the Institutional Animal Care and Use Committee at Arkansas State University and performed in accordance with the Guidelines for the Care and Use of Laboratory Animals. Adult ZSF1 obese (HFpEF) rats at 5 mo (month old) were obtained from Charles River Labs (Wilmington, MA) and used in this study (33, 34). Previous studies have reported that ZSF1 obese rats die at around 12 months of age (35). In our cohort, we found that this group of rats began dying at around 13 months of age. Accordingly, we selected this as the additional (aged) timepoint for the ZSF1 obese group. Importantly, age-matched WT (WKY) rats and ZSF1 lean rats are the widely established and standardized controls for the ZSF1 obese preclinical model. (33, 34, 36). We used both these control groups at both the 5 mo and 13 mo timepoints to maintain consistency with the literature and improve rigor. In addition, we included 20 mo timepoint in both the WT and ZSF1 lean control groups as the oldest aged groups. The total number of mice used were as follows: 5 mo and 13 mo WT control (n=13-19), ZSF1 lean (n=11-35), and ZSF1 obese (n=3-12); 20 mo of WT control (n=8) and ZSF1 lean (n=7).

### Cardiac ultrasound

Echocardiogram was performed to study cardiac function and related physiological assessments including heart rate, interventricular septal dimension in diastole and systole (IVSd and IVSs), left ventricular internal diameter in end-diastole and end-systole (LVIDd and LVIDs), left ventricular posterior wall diameter in diastole and systole (LVPWd and LVPWs). The rats were anesthetized with isoflurane (3% induction and 1.5% maintenance as before) (22, 37) and secured warm on an isothermal pad at 37°C during the procedure following the removal of chest hairs. Two-dimensional echocardiogram measurements were collected from LV short-axis views using a SonoSite M-Turbo system and an ultrasound transducer probe. Raw values were obtained from built-in software and analyzed. Percent fractional shortening (% FS) was calculated using the following formula: [(LVIDd-LVIDs)*100/LVIDd]. Two outliers were eliminated with a z-score greater than 2.

### Terminal isolation

All rats were fully anesthetized with isoflurane anesthesia as before (20, 22) Following assessment of body weights and blood sample collection from the apex of the heart, rats were euthanized using potassium chloride (20 mM; diastolic arrest) injection into the heart. Subsequently, heart, lung, liver, and kidney tissues were dissected and were quickly weighed for morphometric analyses. The tissue samples were flash-frozen in liquid nitrogen and saved at –80°C until use. Left tibias were collected, and lengths were measured for normalization and comparison among the groups.

### Serum thyroid hormone assays

The collected blood was centrifuged at 4000 RPM for 15 minutes at 4°C to isolate the serum. Isolated serum samples were saved at -80°C in small aliquots until further use. Serum total T3 (tT3) and total T4 (tT4) levels were analyzed using respective Accubind enzyme-linked immuno-sorbent assay (ELISA) kits (Monobind Inc., USA) with appropriate controls, following manufacturer instructions. All available mice at all the age timepoints were used for studying both serum tT3 and tT4 ELISA.

### RNA isolation

Left ventricular (LV) tissues (∼60-80 mg) were homogenized in a bullet blender using manufacturer-suggested RINO lysis kits (Next Advances, USA). Total tissue RNA was isolated from homogenized LV tissues using TRIzol reagent (Invitrogen, USA) followed by Invitrogen Purelink RNA kit purification (Invitrogen, USA) following the manufacturer’s recommendations. To ensure the RNA purity, an additional genomic DNA removal step was performed using the RNase-free DNase (Qiagen, USA), and the RNA concentration and quality were initially assessed with Nanodrop.

### Rat lncRNA/mRNA microarray

The total LV RNA from 5 mo groups was studied using the Agilent LncRNA Microarray platform (v2.0; 4 X 44K; ArrayStar Inc., USA) against a comprehensive and robust collection of 13,611 LncRNAs and 24,626 mRNAs. The lncRNAs in the microarray were carefully curated using open-access transcriptomic databases such as RefSeq, Ensembl, and LncRNAdb (Arraystar Inc., USA) (38, 39). Each transcript is represented by a specific exon or splice junction probe for accurate identification. To ensure hybridization quality control, standard positive control (housekeeping genes) and negative control probes were used. RNA quality and quantity were measured by NanoDrop ND-1000 and RNA integrity was assessed by standard denaturing agarose gel electrophoresis (20). The sample preparation and hybridization were performed based on the Agilent One-Color Microarray-Based Gene Expression Analysis protocol with minor modifications. Briefly, samples were amplified and transcribed into fluorescent cRNA along the entire length of the transcripts without 3’ bias utilizing a random priming method (Arraystar Flash RNA Labeling Kit). The labeled cRNAs were purified with RNeasy Mini Kit (Qiagen) and the specific activity and concentration of the fluorescent-labeled cRNAs (pmol Cy3/μg cRNA) were quantified using NanoDrop ND-1000. 1 μg of each labeled cRNA was fragmented in a mix of 10× Blocking Agent and 25× Fragmentation Buffer at 60°C for 30 min, The fragmented cRNAs were diluted in 2× GE Hybridization buffer. A total of 50μl of hybridization solution was dispensed into the gasket slide before assembly to the LncRNA Expression Microarray slide followed by incubation for 17 hours at 65°C in an Agilent Hybridization Oven. Following washing and fixing of the slides, the arrays were scanned using Agilent Scanner G2505C.

To analyze the acquired array images, Agilent Feature Extraction software (version 11.0.1.1) was used. GeneSpring GX v12.1 software (Agilent Technologies) was used for quantile normalization (40) and subsequent data processing. Subsequently, after batch effects were removed using COMBAT, low-intensity lncRNAs and mRNAs were filtered. Gene Set Enrichment Analysis (GSEA) module and GseaPreRanked modules were used to perform GSEA analysis of mRNAs. LncRNAs (at least three samples) were analyzed using the GseaPreRanked module by ranking the Pearson correlation coefficient between the one of selected lncRNAs (10 most differentially expressed in up and 10 in down lncRNAs) and all mRNAs. All arrays passed multiple quality control tests.

The statistically significant differentially expressed lncRNAs and mRNAs were identified through Fold Change filtering between samples and through Volcano filtering between groups. R software was used for Hierarchical Clustering. The thresholds were kept at a fold-change >= 1.5, a P-value <0.05. The fold-change between log2 transformed normalized intensities, *norm1* compared with *norm0* for example, can be calculated as fold-change = 2^|norm1-norm0|^. The initial microarray analyses and data visualization with box plots, scatter plots, volcano plots, and heatmaps contained targets only with p-value<0.05 but were not filtered for false discovery rates. (FDR). Therefore, we further refined the data and presented the scatter plots only with targets satisfying both p<0.05 and FDR<0.05 for improved reliability. For identification of differentially expressed mRNAs with statistical significance, the fold change analysis was performed, and the cut-offs were again set at a fold-change >= 1.5, a P-value <0.05, and an FDR <0.05. Gene Ontology (GO) and Kyoto Encyclopedia of Genes and Genomes (KEGG) pathway analyses were performed on differentially expressed mRNAs. topGO package in the R environment was used for GO enrichment analyses, statistical computing, and graphics. Fisher’s exact test was used for Pathway analysis. The datasets are presented in Supplemental files (1–5).

### Quantitative Real-time PCR

Based on the top hits from the lncRNA microarray, we further investigated the expression patterns of select, significant lncRNA targets (expanded later) at all possible and available timepoints using real-time quantitative PCR (RT-qPCR). First, 1 µg of isolated RNA was converted to cDNA using an RT^2^ first-strand synthesis kit (Qiagen, USA), following manufacturer instructions. qRT-PCR was performed using iTaq universal SYBR Green mastermix (BioRad, USA) in CFX384 (BioRad, USA). All the primers were designed using the PrimerQuest tool (IDT, USA). Normalization was performed with Glyceraldehyde 3-phosphate dehydrogenase (*GAPDH*), Ribosomal Protein Lateral Stalk Subunit P1 (*Rplp1*) and beta 2-microglobulin (*b2m*) genes. Following the removal of low-quality samples with low Cq values and low-quality melting curves, automated data analysis was performed with CFX Maestro software (BioRad, USA) using the default 2^-ΔΔCT method. Fold change of at least ±1.5 and p<0.05 was considered significant. The results were expressed and plotted as relative fold change ± standard error.

### Detection of Altered Inflammatory Factors associated with HFpEF

Both serum and LV tissue lysates from early (5 mo) WT control, ZSF1 lean and ZSF1 obese as well as late (13 mo) ZSF1 obese were used to identify key alterations in inflammatory factors. The tissue lysates were prepared following standard protocol. Briefly, LV tissue sections were homogenized in 300 μL RIPA buffer and Protease Inhibitor cocktails with a bullet blender using manufacturer-suggested RINO lysis kits (Next Advances, USA). The tissue lysates were then agitated at 4C for 2 hours and the supernatants were collected following centrifugation at 16,000 RPM for 20 mins at 4C. The protein quantifications were performed using a Pierce BCA protein kit, following the manufacturer’s guidelines. Subsequently, diluted tissue lysates with a final protein concentration of 25 ug and 1:2 diluted serum samples (with manufacturer-provided sample diluent to a final volume of 100 µL) were used for the streptavidin/biotin-based G-Series Rat Cytokine Array 9 (Ray Biotech, USA), following manufacturer instructions. The slides were dried using compressed Nitrogen gas flow followed by fluorescent imaging and densitometry analyses.

Biotinylated IgG was used as positive control and for signal normalization, detection monitoring, and image orientation. The empty spots on the array were used as negative control or background. The raw data were normalized by RayBiotech analysis using *X(nY) = X(Y) x P1/P(Y),* wherein *P1* refers to the average signal density of the positive control spots on the reference array, *P(Y)* refers to the average signal density of the positive control spots on Array Y, *X(Y)* refers to the signal density for a particular spot on Array for sample “Y”, *X(nY)* refers to the normalized value for that particular spot “X” on Array for sample “Y”. The extracted raw data from quadruplicated assays of 40 distinct inflammatory target proteins were grouped and statistically analyzed in GraphPad Prism using one-way ANOVA with Tukey multiple comparison tests.

### Statistical analyses

The data and statistical analysis for microarray have already been described in section 2.6 above. All results were expressed as means ± standard deviation unless noted otherwise. Statistical data analyses were performed with GraphPad Prism (Versions 9.5.1 and 8.0.2) software. Analyses of groupwise comparisons were carried out using One-way or Two-way analysis of variance (ANOVA) with Tukey’s multiple comparisons tests, or t-test as appropriate. A P-value<0.05 was considered statistically significant.

## Results

### ZSF obese HFpEF rats exhibit preserved contractile function

ZSF1 obese HFpEF rats at 5 mo showed partially increased LV dimensions in systole and diastole compared to both WT and ZSF1 lean controls (Table 1). Nonetheless, the fractional shortening is preserved in all the groups, especially the ZSF1 obese HFpEF rats, suggesting a suitable model for HFpEF (in association with other HF signs discussed later). Our results also corroborate with previously reported findings, which showed a significant decrease in heart rate in 5 mo ZSF1 obese HFpEF males compared to ZSF1 lean counterparts (41) (35). As characterized in other studies (11, 41), preliminary data showed that obese HFpEF rats also showed increased E/E’ indicating diastolic dysfunction (Supplemental Fig S1).

**Table-1:**
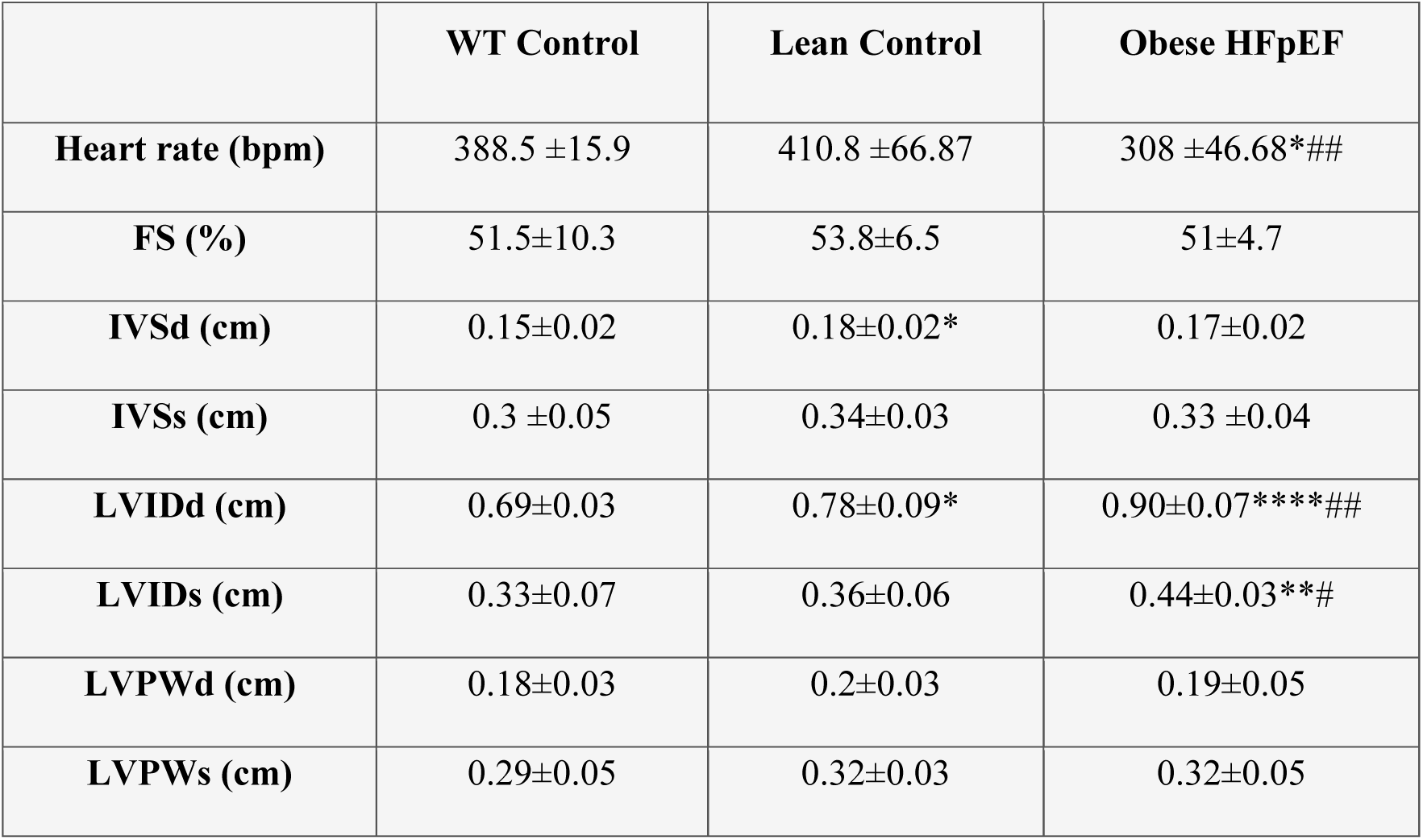
Preserved Ejection Fraction in Male ZSF1 Obese Hearts

### Serum TH levels were significantly decreased in both early and late HFpEF

TH ELISA revealed that total serum T3 (Fig.1A) and T4 (Fig.1B) levels significantly decreased in 5 mo male ZSF1 obese HFpEF rats compared to male ZSF1 lean controls. In addition, serum T4 levels remained significantly low in ZSF1 obese HFpEF rats compared to the ZSF1 lean controls even at the 13 mo timepoint (Fig.1B). In females, serum tT3 levels were significantly decreased in ZSF1 lean controls at both 13 mo and 20 mo timepoints compared to 5 mo (Supplemental Figure. S2A). In males, serum tT3 levels were significantly low in 13 mo vs 5 mo group (Fig. 1C). In addition, based on 2-way ANOVA, there was a significant interaction (p<0.05) between phenotype/genotype and age. Interestingly, tT4 levels did not reveal any significant alterations among WT and ZSF1 lean controls over time in both males (Fig 1D) and females (Supplemental figure S2B). Taken together, the results indicate a significant and persistent hypothyroid phenotype associated with both early and late stages of the HFpEF pathology.

**Figure 1:**
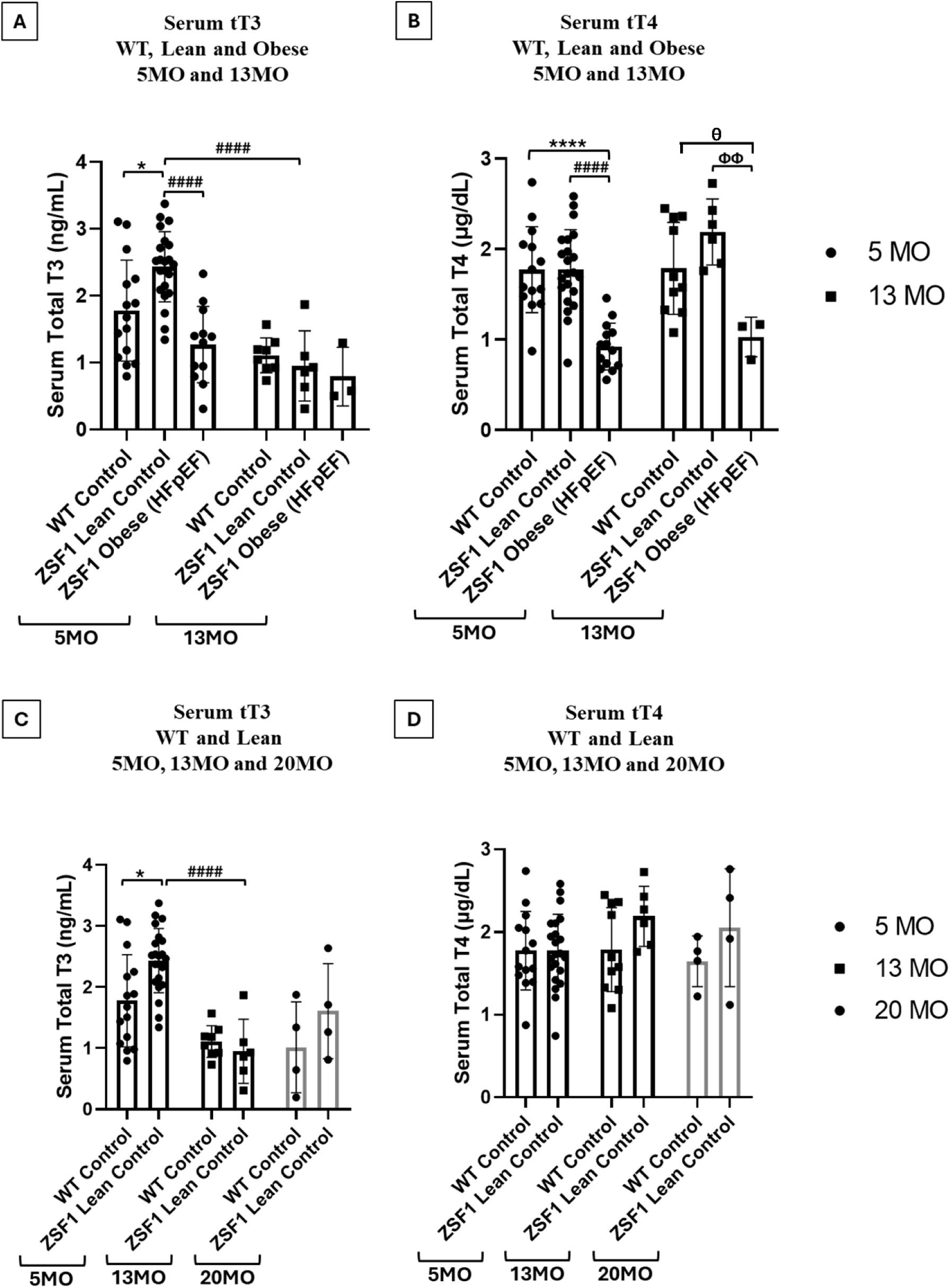
Aging HFpEF males show persistent hypothyroidism: Serum **(A)** total T3 and **(B)** total T4 levels in 5 mo and 13 mo male WT control, ZSF1 lean control, and ZSF1 obese rats; Serum **(C)** total T3 and **(D)** total T4 levels in 5 mo, 13 mo and 20 mo male WT control and ZSF1 lean control. All values are analyzed using Two-way ANOVA (TWA) and presented as means ± standard deviation; WT: Wild Type; T4: Thyroxine; T3: Triiodothyronine; *p<0.05; ****p<0.0001 vs 5 mo WT control (TWA); ^####^p < 0.0001 vs 5 mo ZSF1 lean control (TWA); ^ΦΦ^p<0.01vs 13 mo ZSF1 lean control (TWA); ^θ^p<0.05 vs. 13 mo obese HFpEF (t-test).

### ZSF1 obese HFpEF rats showed hypertrophy of the heart and other systems

Morphometric analyses showed increased heart weight and LV weight indicating cardiac hypertrophy in ZSF1 obese HFpEF rats (Fig.2B-D). Normalizing body and organ weights with tibial length did not show major differences compared to plain weights. Nonetheless, we have shown the tibial length results in the Supplemental Word file. Comparative analyses of 5, 13, and 20 mo WT and ZSF1 lean controls (Fig. 3A-I; Supplemental Figure S5A-D) showed a significant increase in LV weights in both males (Figure 3C) and females (Supplemental Figure S5C) indicating induction of left ventricular hypertrophy in aged ZSF1 lean controls. Since HFpEF is associated with comorbidities and other organs have been shown to be affected in early HFpEF, we studied them in our aged rats (3, 12, 36). Male 5, 13, and 20 mo WT vs ZSF1 lean control comparison also showed significant increases in right lung weight (Fig. 3E), both kidney weights (Fig. 3G-H) and liver weight (Fig. 3I) in ZSF1 lean controls at all timepoints.

**Figure 2:**
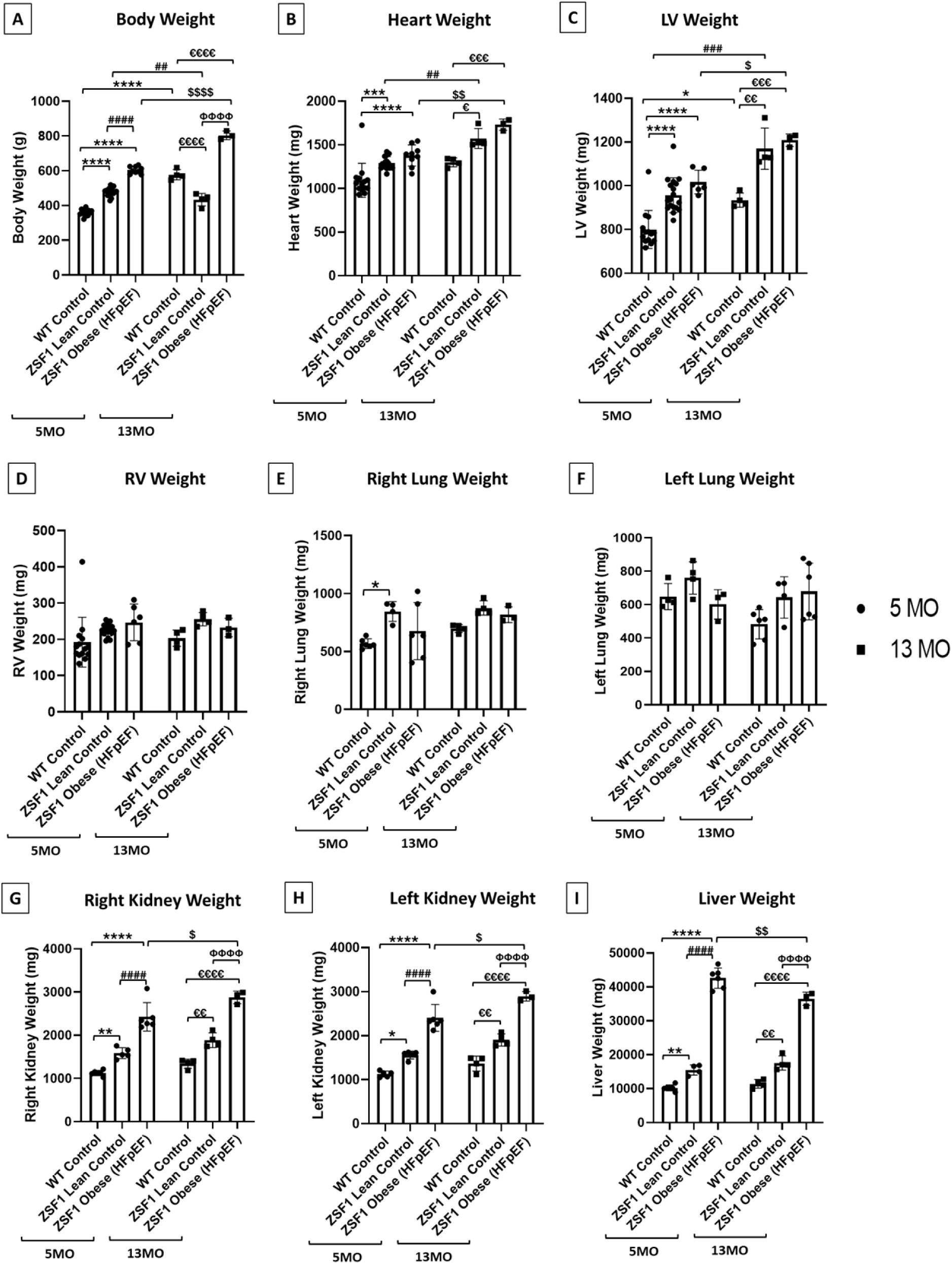
Morphometric analyses. **(Male)** of 5 mo and 13 mo rats from WT control, ZSF1 lean control and ZSF1 obese (HFpEF) groups: **(A)** Body weight; **(B)** Heart weight; **(C)** LV weight; **(D)** RV weight; **(E)** Right lung weight; **(F)** Left lung weight; **(G)** Right kidney weight; **(H)** Left kidney weight and **(I)** Liver weight. All values are analyzed using Two-way ANOVA and presented as means ± standard deviation; TH: thyroid hormone; WT: Wild-type, LV: Left ventricle, RV: Right ventricle, *p<0.05, **p<0.01, ***p<0.001, ****p<0.0001 vs 5 mo WT control; ^##^p<0.01, ^###^p<0.001, ^####^p<0.0001 vs 5 mo ZSF1 lean control; ^$^p<0.05, ^$$^p<0.01, ^$$$$^p<0.0001 vs 5 mo ZSF1 obese (HFpEF); ^ε^p<0.05, ^εε^p<0.01, ^εεε^p<0.001, ^εεεε^p<0.0001 vs 13 mo WT control; ^ΦΦΦΦ^p<0.0001 vs 13 mo ZSF1 lean control.

**Figure 3:**
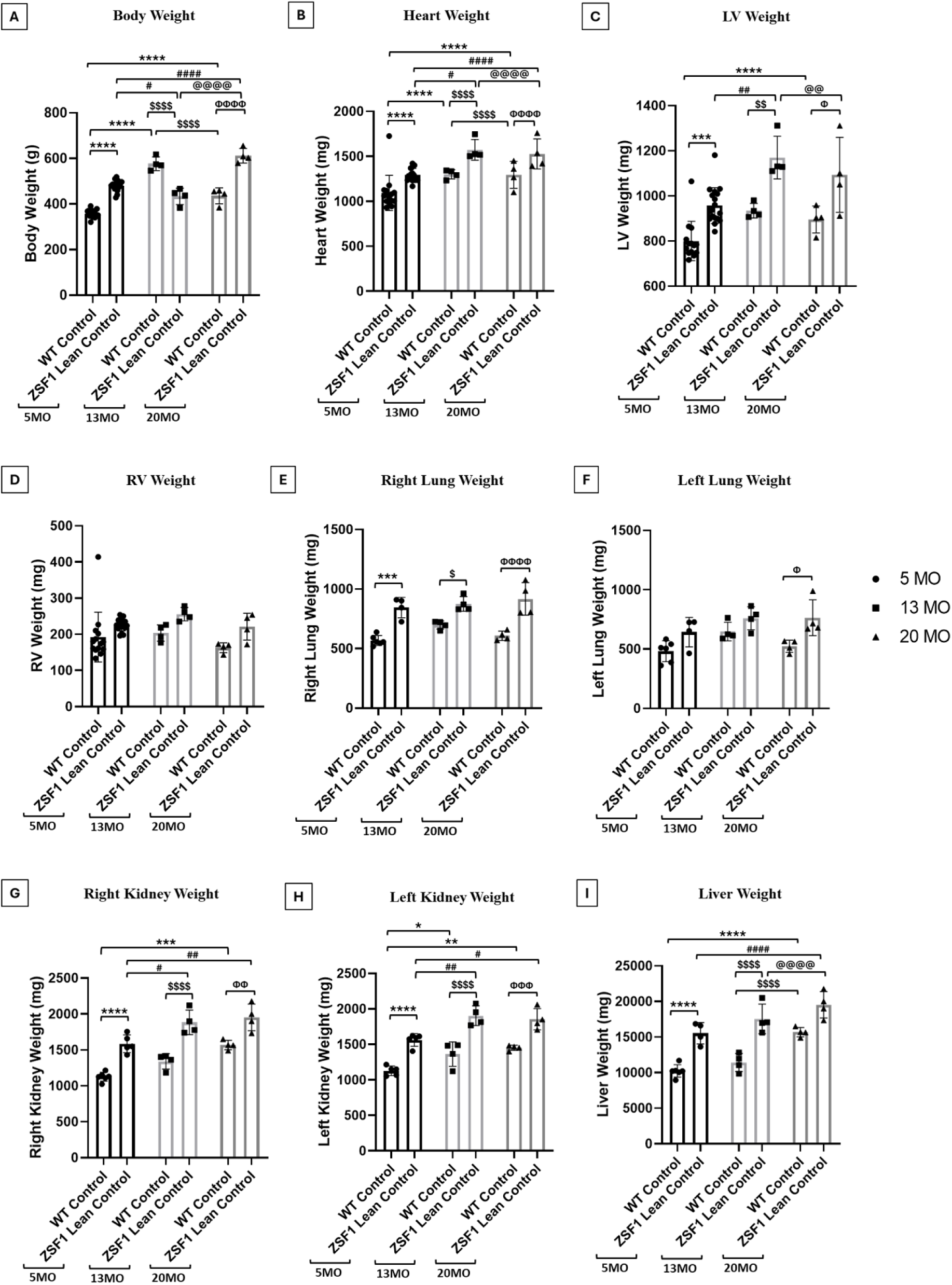
Morphometric analyses. **(Male)** of 5 mo, 13 mo and 20 mo WT control and ZSF1 lean control rats (Male) **(A)** Body weight; **(B)** Heart weight; **(C)** LV weight; **(D)** RV weight; **(E)** Right lung weight; **(F)** Left lung weight; **(G)** Right kidney weight; **(H)** Left kidney weight and **(I)** Liver weight. All values are analyzed using Two-way ANOVA and presented as means ± standard deviation; WT: Wild-type, LV: Left ventricle, RV: Right ventricle weight; *p<0.05, **p<0.01, ***p<0.001, ****p<0.0001 vs 5 mo WT control; ^#^p<0.05, ^##^p<0.01, ^####^p<0.0001 vs 5 mo ZSF1 lean control; ^$^p<0.05, ^$$^p<0.01, ^$$$$^p<0.0001 vs 13 mo WT control; ^@@^p<0.01, ^@@@@^p<0.0001 vs 13 mo ZSF1 lean control; ^Φ^p<0.05, ^ΦΦ^p<0.01, ^ΦΦΦ^p<0.001, ^ΦΦΦΦ^p<0.0001 vs 20 mo WT control.

In the LV, and in both the kidneys, the ZSF1 lean weights were significantly increased at 13 mo compared to the 5 mo group (Fig 3C, G-H). In addition, we have also presented normalized results (by tibial length) for 5 mo and 13 mo WT control, lean control and obese males (Supplemental figure S3A-I), 5 mo, 13 mo and 20 mo WT and lean control males (Supplemental figure S4) and females (Supplemental figure S6). The normalized morphometrics results show similar trends as the organ weight data. Furthermore, 2-way ANOVA analyses of 5 and 13 mo WT control, lean control, and obese HFpEF males showed significant interactions (p<0.001) between phenotype/genotype and age in both body weight and liver weight. Likewise, a comparison of 5, 13, and 20 mo WT and lean control also showed significant interactions (p<0.05) between phenotype/genotype and age in body, heart, and LV weights in both females and males. Additionally, males also showed significant (p<0.01) interaction in liver weight. Together, these findings have thrown new light on hypertrophy of the heart and other related organ systems in this ZSF1 model (42).

### Significant differentially expressed lncRNAs in HFpEF hearts

The analyses showed that several unique lncRNAs were significantly altered (p<0.05; >1.5-fold change with FDR<0.05) in WT control LVs compared to the ZSF1 lean control group or compared to the ZSF1 obese HFpEF group. We have found 29 upregulated (Fig. 4A) and 20 downregulated (Fig. 4B) lncRNAs in the lean control group compared to the WT controls. Furthermore, we have found 57 upregulated (Fig. 4C) 12 and 61 downregulated (Fig. 4D) lncRNAs in ZSF1 obese HFpEF vs WT control comparison. In addition, 12 upregulated and 12 downregulated unique lncRNAs were also found to be common between 5 mo ZSF1 lean vs WT control groups and obese HFpEF vs WT control groups (Supplemental File 1, marked in red). We have also presented additional details such as seqname (sequence identifier), GeneSymbol, RNA length, chromosome number, strand, txStart (transcription start site), txEnd (transcription end site), EntrezID, noncoding RNA characteristic, information about any associated genes, etc.

**Figure 4:**
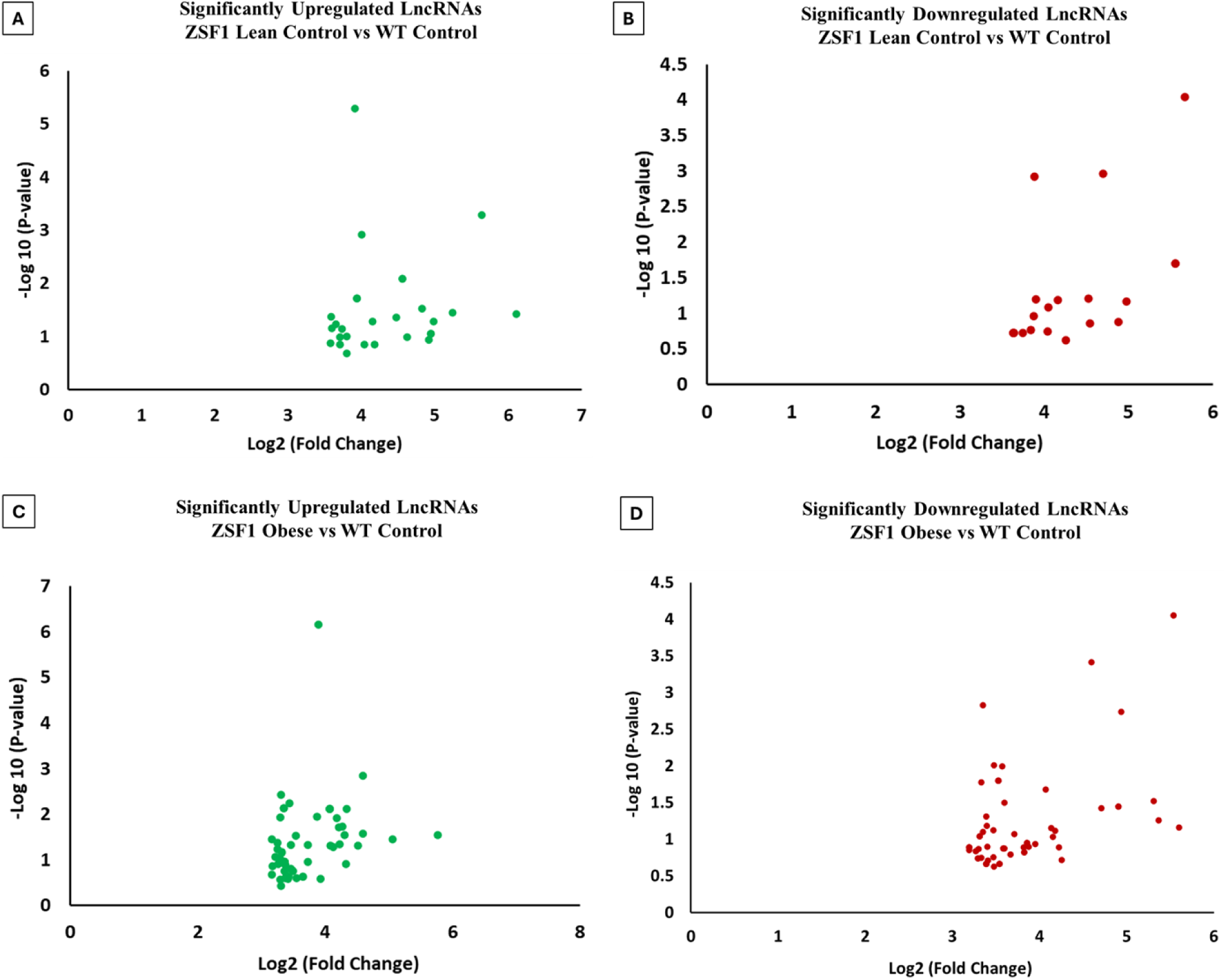
Differentially expressed lncRNAs in HFpEF: LncRNA microarray analyses (p<0.05) and further refinement (fold-change>1.5 and FDR<0.05) of differentially expressed lncRNAs in 5 mo WT control, ZSF1 lean control, and ZSF1 obese rats: **(A)** Upregulated lncRNAs in ZSF1 lean vs WT controls; **(B)** Downregulated lncRNAs in ZSF1 lean vs WT controls; **(C)** Upregulated lncRNAs in ZSF1 obese (HFpEF) vs WT controls; **(D)** Downregulated lncRNAs in ZSF1 obese (HFpEF) vs WT controls.

## Left ventricular mRNA microarray analyses

### Significant differentially expressed mRNAs in HFpEF hearts

mRNA microarray analysis showed numerous mRNAs that were differentially regulated (at least 1.5-fold with P-value and FDR <0.05) (Supplemental File 2) in ZSF1 groups in comparison with WT controls. The ZSF1 lean vs WT control comparison identified 43 upregulated (Figure 5C) and 33 downregulated (Figure 5D) mRNA transcripts. On the other hand, ZSF1 obese HFpEF and WT Control showed 119 upregulated (Figure 5E) and 122 downregulated (Figure 5F) mRNAs. Differential expression analysis in ZSF1 lean and ZSF1 obese HFpEF comparison showed only 1 upregulated (*Sik1*) (Figure 5A) and 1 downregulated (*Anxa13*) (Figure 5B) mRNA transcripts.

**Figure 5:**
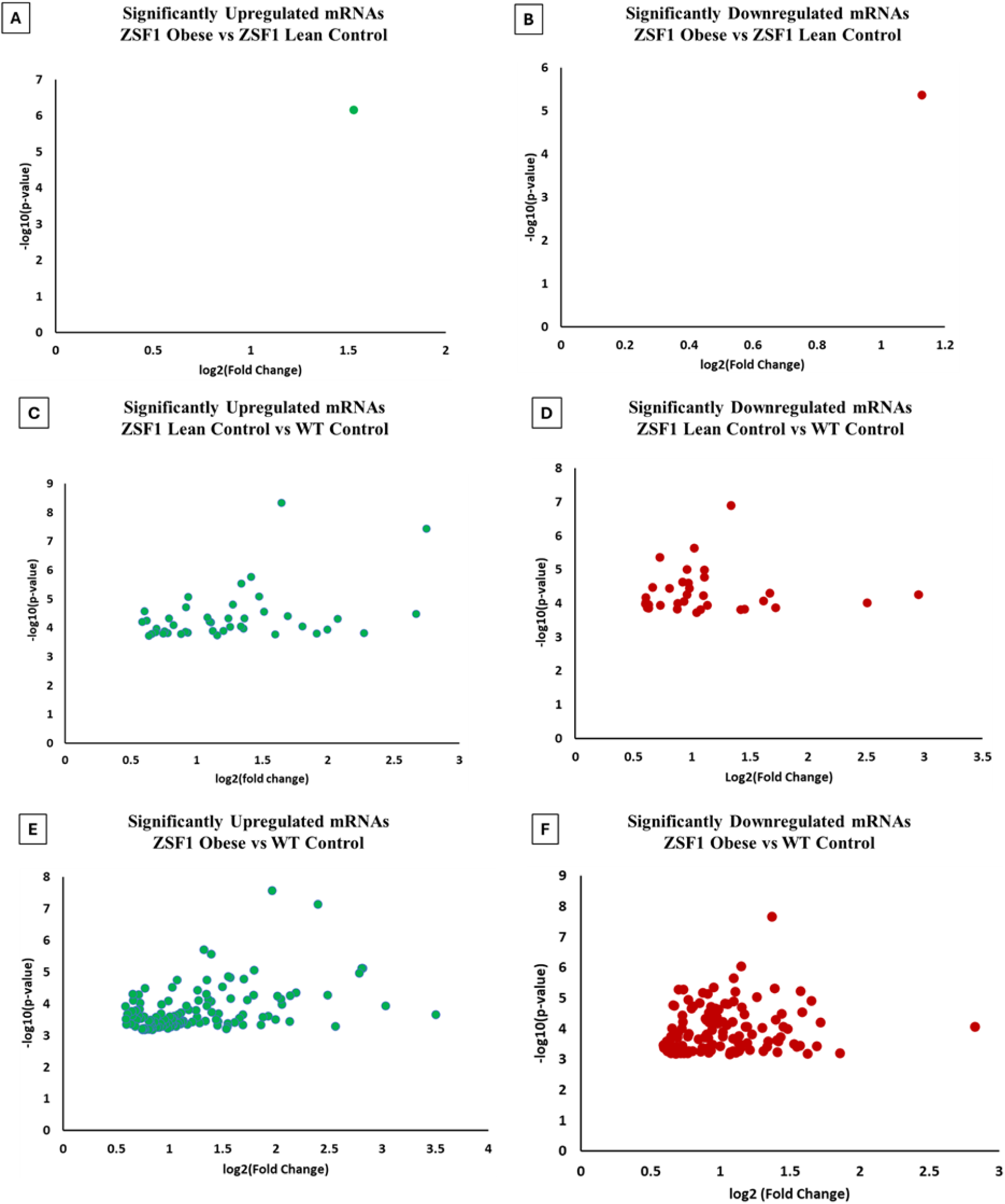
Differentially expressed mRNAs in HFpEF: mRNA microarray analyses (p<0.05) and further refinement (fold-change>1.5 and FDR<0.05) identified differentially expressed mRNAs in 5 mo WT control, ZSF1 lean controland ZSF1 obese rats: **(A)** Upregulated mRNAs in ZSF1 obese (HFpEF) vs lean controls; **(B)** Downregulated mRNAs in ZSF1 obese (HFpEF) vs lean controls; **(C)** Upregulated mRNAs in ZSF1 lean vs WT controls; **(D)** Downregulated mRNAs in ZSF1 lean vs WT controls; **(E)** Upregulated mRNAs in ZSF1 obese (HFpEF) vs WT controls; **(F)** Downregulated mRNAs in ZSF1 obese (HFpEF) vs WT control.

In addition, among the up- and down-regulated mRNAs in ZSF1 lean vs WT control comparisons, 26 upregulated and 21 downregulated unique mRNA transcripts were also up- or down-regulated in ZSF1 obese HFpEF vs WT control comparisons (p<0.05; FDR<0.05) (Supplemental File 2). This indicates that several genes from the non-HFpEF (lean) phenotype continue to be involved in the HFpEF (obese) phenotype.

## mRNA enrichment analyses

### GO enrichment analyses

In GO enrichment analyses (Supplemental File 3), lean control vs WT control showed 4 Biological Process (BP) pathway enrichments and 3 Cellular Component (CC) enrichments among the significantly upregulated genes (p<0.05) (Fig. 6B). Within the downregulated genes in lean control vs WT control, we found 8 BP, 1 CC and 1 Molecular Function (MP) enrichments (p<0.05) (Fig. 6C). Groupwise comparison of ZSF1 obese HFpEF and WT control showed 21 GO enriched BPs in significantly upregulated coding genes (Fig. 6A). Interestingly, a closer look of the BP enrichments showed distinct pathway clusters among the two aforesaid comparisons. BP enrichment of significantly upregulated mRNAs in obese HFpEF vs WT control comparison is predominantly a cluster of metabolic pathway genes, whereas BP enrichment of upregulated mRNAs in lean control vs WT control is a cluster of only immune response pathways. Furthermore, BP enrichment analyses of significantly downregulated mRNAs in lean control vs WT control showed a combination of oxygen/gas transport, metabolic pathways, immune response pathways, and others.

**Figure 6:**
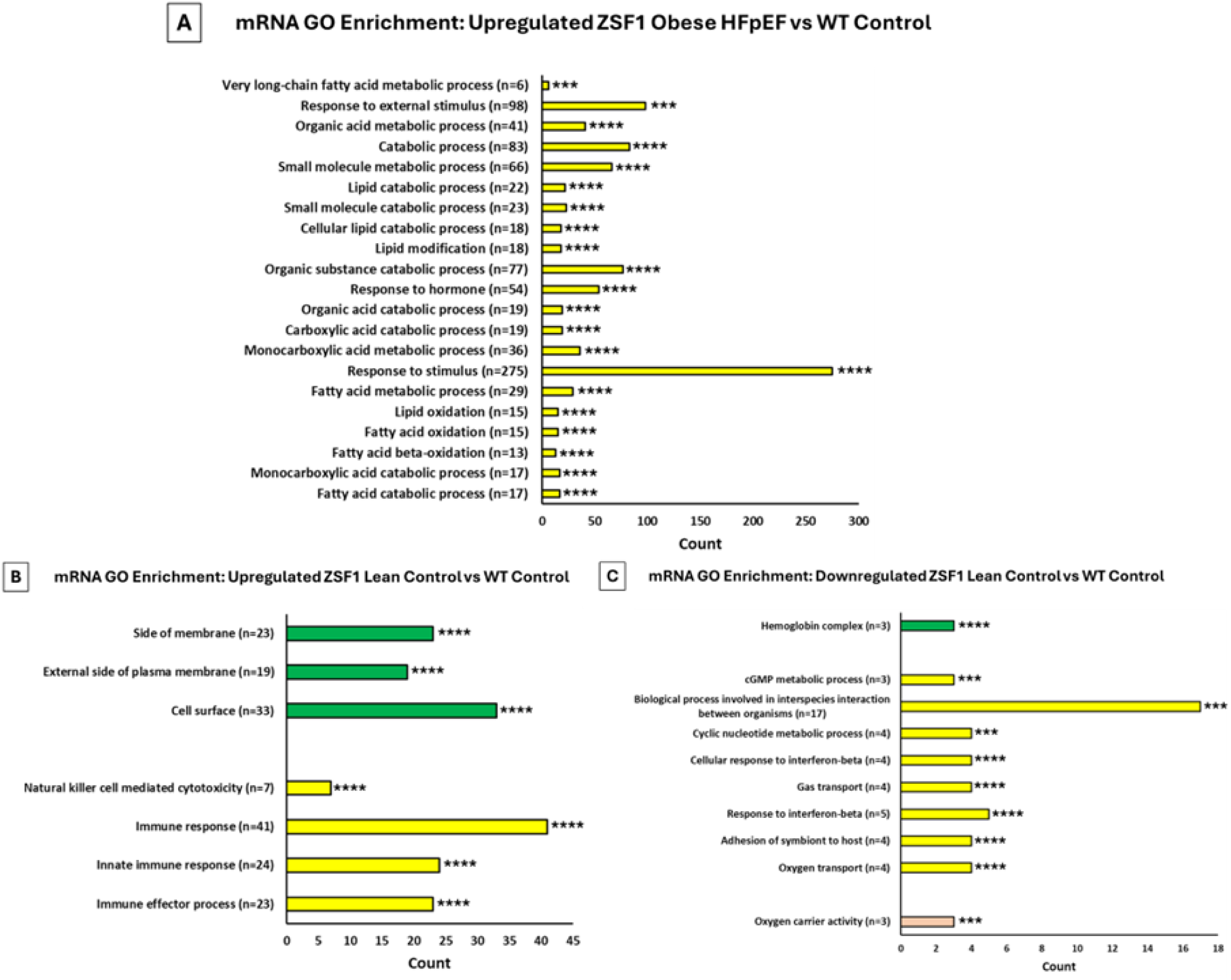
GO enrichment analyses of mRNAs: **(A-C)** Significantly enriched GO terms (p<0.05 and FDR<0.05) in **(A)** Upregulated in ZSF1 obese vs WT control; **(B)** Upregulated in ZSF1 lean vs WT control; **(C)** Downregulated in ZSF1 lean control vs WT control; GO: Gene Ontology; ***p<0.001, ****p<0.0001.

### KEGG pathway enrichment analyses

As listed in Supplemental File 4, the Lean vs WT control comparison yielded two vector-borne disease pathways – Malaria and African trypanosomiasis (Fig. 7C; p<0.05). The obese HFpEF showed upregulation of three different fatty acid pathways – fatty acid elongation, fatty acid metabolism, and biosynthesis of unsaturated fatty acids compared to WT controls (Fig 7D; p<0.05). On the other hand, the estrogen receptor pathway and circadian entrainment pathway were significantly downregulated in the obese HFpEF vs lean control comparison (Fig. 7B; p<0.05). This obese HFpEF vs lean control comparison also showed upregulation of 5 pathways including fatty acid elongation, biosynthesis of unsaturated fatty acids, fatty acid degradation, fatty acid metabolism, and lysine degradation (Fig. 7A; p<0.05).

**Figure 7:**
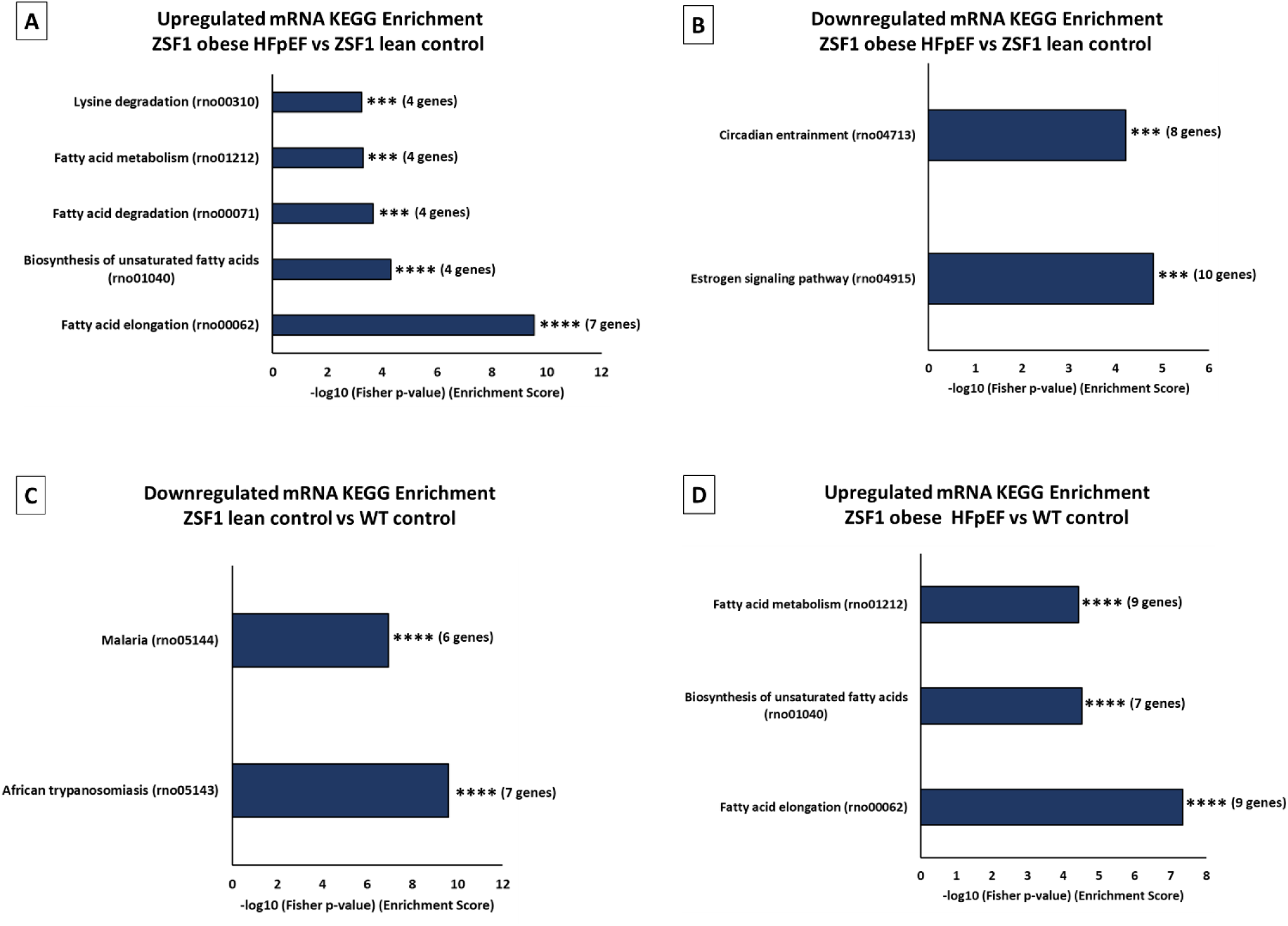
KEGG Enrichment analyses of mRNAs: **(A-D)** Significantly enriched KEGG pathways (p<0.05 and FDR<0.05) in **(A)** Upregulated in ZSF1 obese vs lean control; **(B)** Downregulated in ZSF1 obese vs lean control; **(C)** Downregulated in ZSF1 lean control vs WT control; **(D)** Upregulated in ZSF1 obese vs WT control; KEGG: Kyoto Encyclopedia of Genes and Genomes; ***p<0.001, ****p<0.0001.

### HFpEF hearts showed lncRNA-mRNA association

Similar to results from the above differentially expressed lncRNAs, further analyses showed that several of them had associated mRNAs (coding gene pairs) with a p<0.05. However, further sorting with an FDR<0.05 (Supplemental File 5), showed two potential lncRNA-mRNA associations in the ZSF1 obese HFpEF vs WT control group comparison. None of the other groups showed any significant lncRNA-mRNA associations. *LOC102548445* and *LOC102556567*, two significantly upregulated lincRNAs, were found to be closely associated with 2 mRNA genes respectively – *Ifi44l* and *AABR06107100.1*. While *Ifi44l* was downregulated, the X-chromosomal gene *AABR06107100.1* (likely associated with *Nxf5*) was upregulated in our mRNA DE analyses of obese HFpEF vs WT control (Supplemental File 5). We have also investigated the antisense lncRNAs and associated coding gene pairs, but no groupwise comparisons showed any significantly altered antisense lncRNA-mRNA pairs, after sorting (p-value<0.05, FDR<0.05).

### Real-time quantitative PCR identified unique lncRNAs in older age phenotypes

As discussed earlier, preclinical HFpEF studies commonly focus on the early HFpEF timepoint (∼5 mo). As aging is strongly correlated with the development of HFpEF (42), we aimed to detect whether the lncRNA expression patterns change with age/severity of HFpEF pathologies. In order to identify aging HFpEF-associated lncRNAs, we performed RT-qPCR with select lncRNA targets from the microarray results on samples from all possible and available time points. To ensure high rigor in lncRNA microarray results, we did not include those targets that showed FDR>0.05 (even though p<0.05). We chose the top 3-4 targets from microarray analyses based on the highest fold change (p<0.05 and FDR<0.05) and excluded any genes suspected to be protein-coding or pseudogene. The analyses showed that 13 mo ZSF1 obese HFpEF rats had significantly higher expression levels of cardiac *LOC102555623* (Fig 8A), *LOC102553944* (Fig 8B), *LOC100912003* (Fig 8C) lncRNAs compared to the ZSF1 lean 13 mo group. The expression levels of both *LOC102555623* and *LOC100912003* were also found to be significantly increased in the 13 mo ZSF1 obese group compared to both the ZSF1 obese 5 mo and WT 13 mo groups. Interestingly, the 20 mo ZSF1 lean LVs had a similar level of *LOC102555623* expression compared to the 13 mo ZSF1 obese group. Furthermore, both *LOC102555623* and *LOC102553944* lncRNA expression levels were significantly increased in the ZSF1 lean 20 mo group compared to, WT 20 mo and ZSF1 lean 13 mo, but not when compared to obese HFpEF 13 mo. Additionally, the expression levels of these three lncRNAs showed similar trends among groups in microarray and RT-qPCR at 5 mo. These indicate that the late-stage aged ZSF1 lean 20 mo old rats develop molecular changes similar to the HFpEF group.

**Figure 8.**
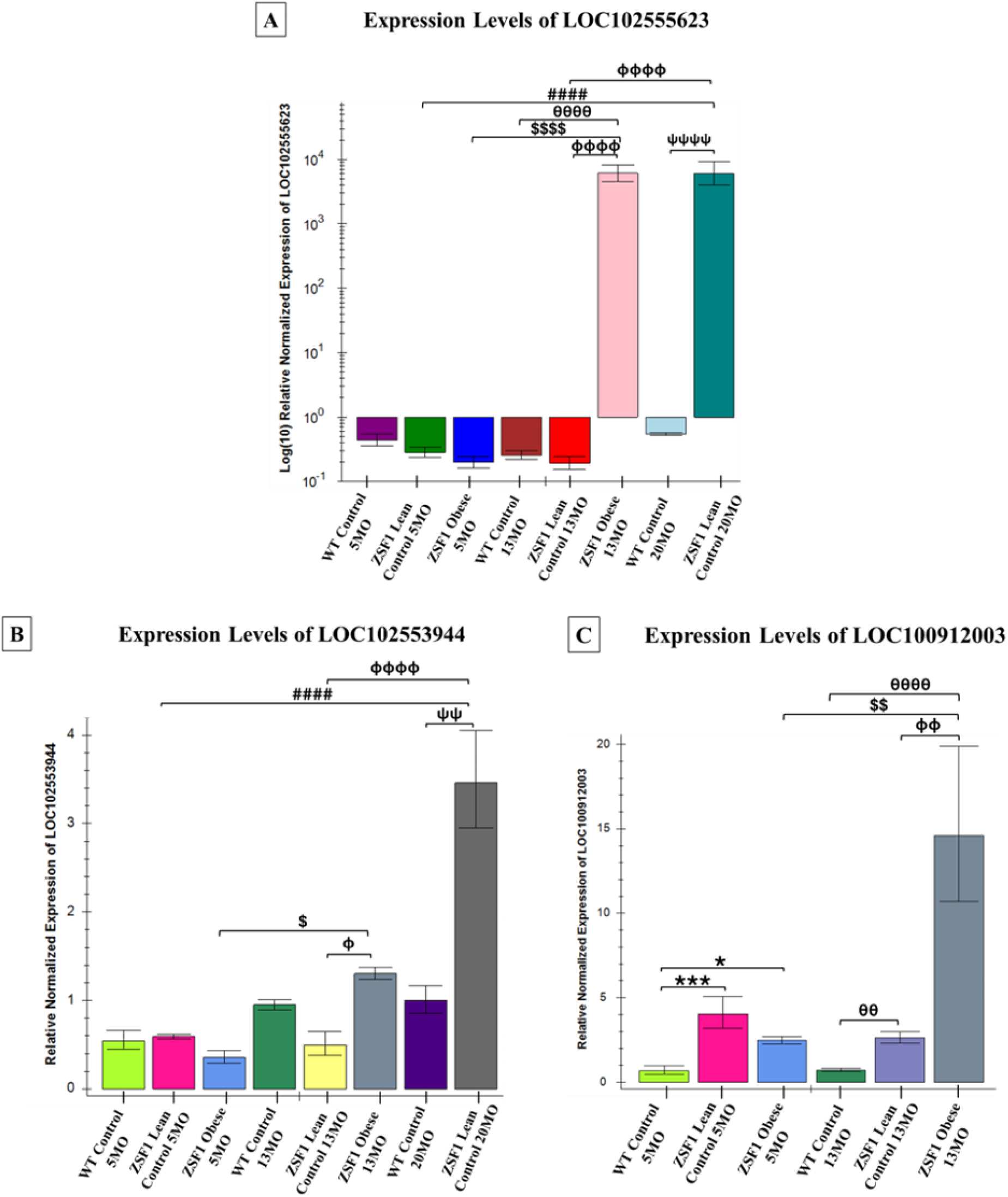
RT-qPCR analyses identified novel lncRNAs associated with aging hypothyroid obese HFpEF. **(A-C)** RT-qPCR of 5 mo, 13 mo +/- 20 mo WT control, ZSF1 lean control and ZSF1 obese HFpEF left ventricular (LV) samples showed significantly altered expression levels of **(A)** *LOC102555623*; **(B)** *LOC102553944* and **(C)** *LOC100912003*. Results are presented as Mean ± Standard error. *p<0.05, ***p<0.001 vs 5 mo WT control; ^####^p<0.0001 vs 5 mo ZSF1 lean control; ^$^p<0.05, ^$$^p<0.01, ^$$$$^p<0.0001 vs 5 mo ZSF1 obese HFpEF; ^θθ^p<0.01, ^θθθθ^p<0.0001 vs 13 mo WT control; ^Φ^p<0.05, ^ΦΦ^p<0.01, ^ΦΦΦΦ^p<0.0001 vs 13 mo ZSF1 lean control; ^ΨΨ^p<0.01, ^ΨΨΨΨ^p<0.0001 vs 20 mo WT control.

We identified one significantly upregulated (*Sik1*) and one downregulated mRNA (*Anxa13*) among the 5 mo lean control vs obese comparisons in the microarray analyses, RT-qPCR showed no significant changes during the late stage of the disease. This indicates the possibility that these genes may be critical during the onset of HFpEF rather than the progression or severity of the disease.

### Aging HFpEF shows altered serum and cardiac inflammatory markers

Chronic low-grade inflammation has been previously reported to be associated with HFpEF (43). Previous work from our lab (20) showed dysregulation of inflammatory pathway genes under altered TH levels. As hypothyroidism is a characteristic feature of HFpEF (44), we tried to identify clinically relevant inflammatory cytokine markers associated with the early and late stages of HFpEF. Moreover, significant alterations in inflammatory pathway genes in our GO analyses primed our interest to further investigate their protein profiles. Inflammation arrays in 5 mo WT Control, ZSF1 lean control and ZSF1 obese HFpEF, and 13 mo ZSF1 obese HFpEF rats showed markedly different trends in inflammatory marker levels in both LV tissue lysates and in serum (Figure 9A-9L; p<0.05). A majority of the alterations in inflammatory markers were observed in the LV tissue lysates.

**Figure 9:**
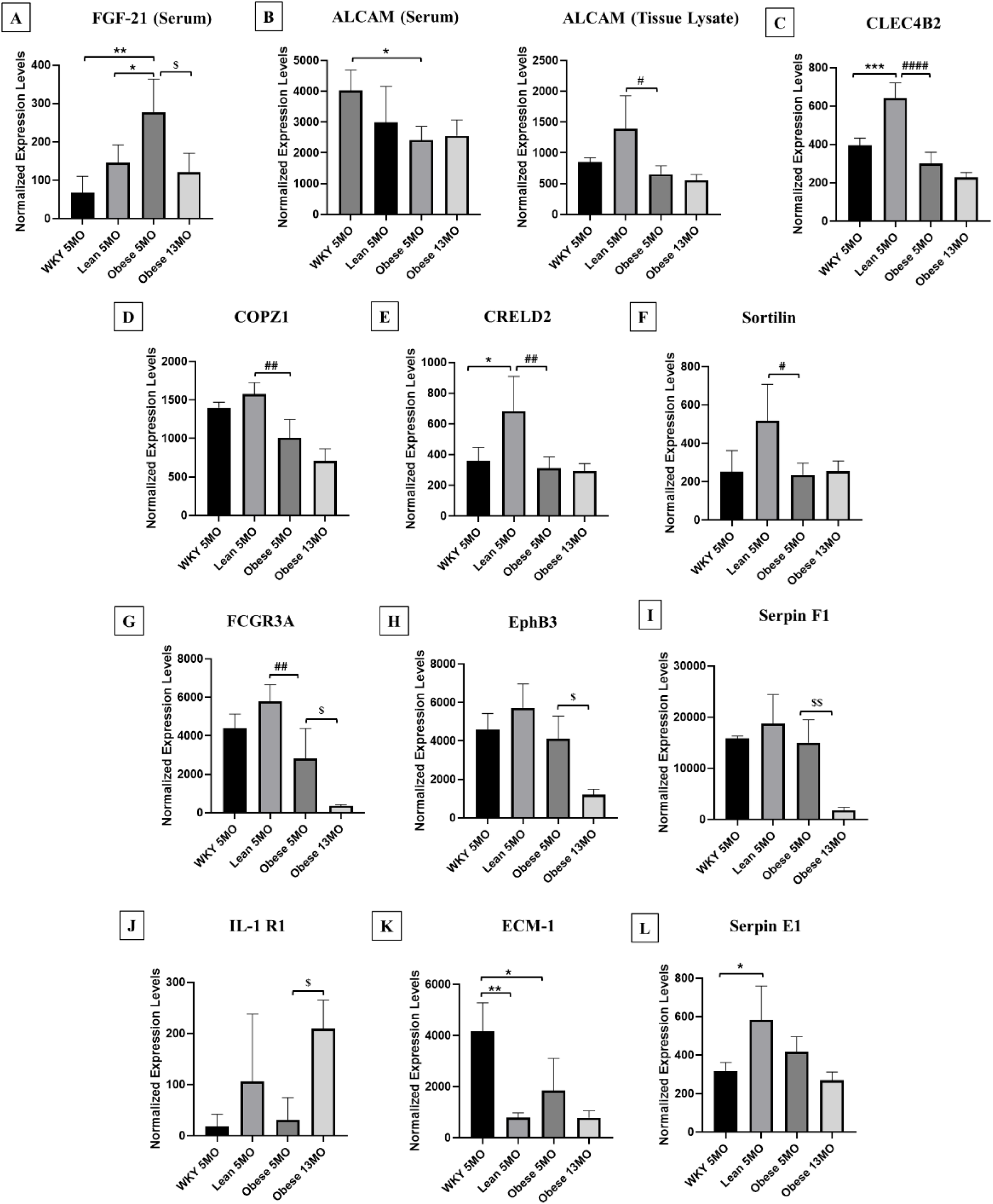
Aging HFpEF shows Altered Serum and Cardiac Tissue Inflammatory Profiles: Serum and cardiac tissue inflammatory factor profiles in 5 mo WT control, 5 mo ZSF1 lean control, 5 mo ZSF1 obese HFpEF and 13 mo ZSF1 obese HFpEF: **(A)** FGF-21 (Serum); **(B)** ALCAM (Left subset: Serum; Right subset: Tissue Lysate); **(C)** CLEC4B2 (Tissue); **(D)** COPZ1 (Tissue); **(E)** CRELD2 (Tissue); **(F)** Sortilin (Tissue); **(G)** FCGR3A (Tissue); **(H)** EphB3 (Tissue) **(I)** Serpin F1 (Tissue); **(J)** IL-1 R1 (Tissue); **(K)** ECM-1(Tissue); **(L)** Serpin E1 (Tissue). All values are presented as means ± standard deviation. *p<0.05, **p<0.01, ***p<0.001 vs. 5 mo WT control; ^#^p<0.05, ^##^p<0.01, ^####^p<0.0001 vs. 5 mo ZSF1 lean control; ^$^p<0.05, ^$$^p<0.01 vs. 5 mo ZSF1 obese HFpEF.

Compared to lean rats, 5 mo (early) HFpEF showed a significant increase in serum FGF-21 levels. This indicates that serum FGF-21 may be used as a potential biomarker to detect early onset of HFpEF (discussed later). LV tissue ALCAM, CLEC4B2, COPZ1, CRELD2, and Sortilin levels were significantly downregulated in 5 mo obese hearts compared to the lean controls and remained low in the 13 mo obese rats. Serum ALCAM levels showed a significant decrease in the 5 mo obese rats compared to age-matched WT Controls. FCGR3A, not only showed a significant reduction in 5 mo obese LVs compared to the 5 mo lean counterparts (similar to the tissue FGF21) but also further significantly reduced in the 13 mo obese HFpEF group. Interestingly, the tissue levels of EphB3 and Serpin F1 were not significantly decreased in obese rats at the 5 mo time point compared to the controls but reduced significantly in late (13 mo) obese hearts compared to the 5 mo obese group. This indicates that EphB3 and Serpin F1 have the potential to serve as inflammatory markers to identify the severity of HFpEF. On the contrary, while tissue levels of IL-1 R1 did not show any significant difference between 5 mo lean vs 5 mo obese comparisons, it was significantly increased in 13 mo obese rats compared to the 5 mo obese rats. This indicates that similar to EphB3 and Serpin F1, IL-1 R1 may also serve as a tissue biomarker to detect the severity of HFpEF. We found that ECM-1 levels were significantly reduced in both 5 mo lean controls and obese rats indicating that it may serve as a biomarker for hypertension. Furthermore, similar to CLEC4B2 and CRELD2, Serpin E1 levels were significantly increased in 5 mo lean controls compared to 5 mo WT controls. These indicate the potential for the development of multiple biomarkers for HFpEF and/or hypertension.

## Discussion

This is the first preclinical study to our knowledge investigating the later stages of HFpEF. Compared to the lean controls, we uncovered that serum T4 levels were not only significantly decreased in the 5 mo obese HFpEF rats, but also remained low in the 13 mo late HFpEF stage. This model provides a unique opportunity to shed light on possible mechanisms seen in human HFpEF (44). This is also the first known report that has studied the effects of the ZSF1 phenotype for the longest possible duration (20 months). The role of lncRNAs in HFpEF as a geriatric syndrome with thyroid dysfunction has been unclear (45). Five months is the earliest timepoint where HFpEF is commonly studied. In support of detecting a broad array of targets that may help with preventive screening (in future studies) of potential biomarkers associated with aged hypothyroid HFpEF, we conducted the microarray at 5 mo timepoint and compared them with select top hit targets via RT-qPCR at the 13 mo timepoint. Accordingly, using lncRNA-specific microarray and subsequent RT-qPCR analyses, we discovered three novel lncRNAs that have not been shown before. *LOC102555623, LOC102553944, and LOC100912003* were significantly elevated in the 13 mo obese HFpEF LVs compared to their 5 mo obese counterparts. In addition, although all three lncRNAs were not significantly altered in the 5 mo obese groups compared to the 5 lean groups, all three were significantly higher in 13 mo obese groups compared to 13 lean groups. Furthermore, our findings that the levels of these lncRNAs are either equally higher or much higher in the 20 mo ZSF1 lean rats (compared to the 13 mo ZSF1 obese HFpEF rats) suggest the former may be approaching HF phenotype at this time-point.

Quantitative trait loci (QTL) (46) search in the Rat Genome Database showed that *LOC102553944* is significantly associated with multiple relevant traits including body weight gain, feed conversion ratio, plasma corticosterone level, plasma insulin level, pancreas wet weight, and plasma free fatty acids level. In addition, *LOC102555623* was associated with systolic blood pressure, retroperitoneal fat pad weight to body weight ratio, abdominal subcutaneous fat pad weight, lean tissue morphological measurement, abdominal fat pad weight to body weight ratio, epididymal fat pad weight to body weight ratio, body weight gain, both kidneys wet weight to body weight ratio. Further investigation can be helpful in understanding the pathophysiological roles, diagnostic potential, and druggability of these three lncRNAs for aging hypothyroid HFpEF. LncRNAs often exhibit a high level of structural conservation along with conserved functions across species (47). Thus, our results suggest that lncRNAs may play an important role in the progression and/or severity of aging HFpEF. We have also identified one significantly upregulated Salt inducible kinase 1 (*Sik1*) and one downregulated Annexin A13 (*Anxa13*) mRNA in early (5 mo) ZSF1 lean control vs obese HFpEF comparison in the microarray analyses. *Sik1* has been previously shown to be involved in promoting pathologic cardiac remodeling but has not been implicated in HFpEF (48). On the other hand, *Anxa13* has never been reported to be conclusively involved in the development or progression of CVDs. Taken together, this is the first report showing dysregulation of both *Sik1* and *Anxa13* in HFpEF.

Hypertension is one of the most common co-morbidities in HFpEF patients. It is not only a modifiable risk factor but has also been found to be a critical contributor to the pathogenesis and prognosis of the disease (9). Both ZSF1 lean controls and ZSF1 obese rats are hypertensive at about 5 mo age or earlier (41, 49, 50), and we included WT controls to identify potential molecular factors influencing the genotypes. Interestingly, we found that 26 upregulated and 21 downregulated mRNA genes were common between two comparisons--5 mo WT control vs ZSF1 lean comparison and 5 mo WT control vs obese comparison (Supplemental File 3, marked in red). Similarly, 12 upregulated and 12 downregulated lncRNA genes were also found to be common between the same two aforesaid comparisons. It is possible that these genes may be associated with hypertension, an important common feature among both ZSF1 lean and obese rats. The commonalities in these noncoding and coding RNAs between both non-HF (lean) and HF (obese) phenotypes suggest that drugs targeting these genes may help improve both these pathological conditions. Compared to WT control, obese hearts showed GO enrichment changes predominantly in metabolic and lipid pathways. On the other hand, the lean control GO pathways showed changes predominantly in immune mechanisms. Similar findings were found in the KEGG analyses. Compared to both the control groups, the LVs in the obese HFpEF group showed significant upregulation predominantly in lipid-based metabolic pathways. These findings suggest that metabolic/lipid factors may play a larger role in the HFpEF hearts alongside other comorbidities such as hypertension, diabetes mellitus, etc. These findings may also help in identifying nodes and targets in the development of novel interventions for HFpEF.

The comorbidities seen in HFpEF are considered to be secondary to systemic inflammation resulting in abnormal levels of inflammatory cytokines. In particular, metabolic inflammation or metainflammation is considered central to HFpEF pathophysiology (51, 52). The ZSF1 model serves as one of the best tools that embodies the clinical manifestations of HFpEF. We recently showed that multiple inflammatory and immune-related lncRNAs were significantly impaired in the LVs of experimentally induced hypothyroid cardiac dysfunction (20). Our current study revealed that this is also observed in the hypothyroid HFpEF LVs, albeit via a different set of aforesaid lncRNAs. Inflammatory biomarkers, including C-reactive protein (CRP), interleukin (IL)-1, tumor necrosis factor (TNF)-α (43) and its receptors, TNFR1 and TNFR2 (53), IL-6 the chemokine (C–C motif) ligand 2 (CCL2) (54), etc. are often elevated in patients with HFpEF. Chronic, low-grade, systemic inflammation might have detrimental effects on myocardial structure and function. In agreement with the previously reported association of inflammation in HFpEF, our microarray analyses of the 5 mo HFpEF group also showed multiple, significantly enriched inflammatory pathways in the mRNA GO analyses. This primed our interest to identify previously unreported inflammatory proteins in aging hypothyroid HFpEF.

FGF-21 levels were significantly increased in the serum of the early obese HFpEF rats compared to the lean controls. This concurs with the clinical findings that circulating FGF21 is increased in HFpEF (55, 56). It would be valuable to further investigate its role in possible inflammaging mechanisms (57). We also found multiple targets significantly altered in late vs early HFpEF LV tissues. While FCGR3A (CD16), EphB3, and Serpin F1 levels decreased in 13 mo obese HFpEF compared to 5 mo obese HFpEF, IL1-R1 levels were found to be significantly increased in the 13 mo obese HFpEF compared to the 5 mo counterparts. Although IL1-R1 has been implicated in aging-associated cardiomyopathy (58), FCGR3A (CD16), EphB3, and Serpin F1 have not been shown. This is the first report that uncovers the importance of these molecules in aging hypothyroid HFpEF. Moreover, given the significant downregulation of ECM-1 in all the ZSF1 groups and time-points studied in this inflammation assay, it is possible that targeting this gene may be helpful to overcome both hypertension and HFpEF (early and late) or hypertension alone (reported in both the genotypes). The potential interactions among the 3 lncRNAs, the 2 mRNAs, and the panel of inflammation-related markers that we detected are not clear and remain a ripe area for future investigations.

The limitations of the study include the following. Although we used both females and males in the majority of the groups, we could only acquire males for the ZSF1 obese group of rats. Future assessment of ZSF1 obese female rats at both 5 mo and 13 mo can help understand any sex-related differences in HFpEF development and severity. Although this study does not establish a strong causative relationship, it does demonstrate that hypothyroidism is an important comorbid factor in HFpEF and may contribute/worsen the pathology in association with noncoding RNA and inflammatory mechanisms. Since hypothyroidism was observed not only in the old HFpEF groups, but also in our young HFpEF groups, it is likely that the hypothyroidism is not primarily due to aging, but rather a key factor in the HFpEF development. It would also be useful for future studies to employ next-generation RNA sequencing to identify any transcripts that may have been missed by microarray.

In summary, we have shown long-term mechanisms in aged HFpEF and hypertensive heart disease. More specifically, we have identified novel cardiac lncRNAs and inflammatory targets associated with persistent hypothyroidism in the aging HFpEF syndrome. Treatment options for HFpEF are limited and future studies can help towards further understanding of mechanisms and development of better diagnostic and therapeutic targets for this life-threatening condition that presents with severe morbidity and mortality.

## Declarations

### Ethics approval and consent to participate

Not applicable.

### Consent for publication

All authors approved the final manuscript before submission.

### Availability of data and materials

Not applicable.

### Competing interests

The authors declare no competing interests.

### Funding

We thank the funding sources to Dr. Viswanathan Rajagopalan from the New York Institute of Technology College of Osteopathic Medicine at Arkansas State University and Arkansas Biosciences Institute (the major research component of the Arkansas Tobacco Settlement Proceeds Act of 2000). We also thank the American Physiological Society for the Summer Undergraduate Research Fellowship. The funders had no role in study design, data collection and analysis, decision to publish, or preparation of the manuscript.

### Author Contributions

V.R.: Conceptualization, Methodology, Validation, Formal Analysis, Investigation, Resources, Writing— Original draft preparation, Writing—Reviewing and Editing, Visualization, Supervision, Project administration, Funding acquisition. S.C.: Methodology, Validation, Formal Analysis, Investigation, Writing—Original draft preparation, Writing—Reviewing and Editing, Visualization. CB, BD, SL, CH: Methodology, Validation, Formal Analysis, Investigation. All authors have read and agreed to the published version of the manuscript.

## Supporting information

Supplemental Results

## Acknowledgments

We acknowledge multiple student trainees who learned techniques in the laboratory. We would also like to acknowledge Dr. Ruhi Shethwala for her involvement in the study.

## Notes

### Competing Interest Statement

The authors have declared no competing interest.

