## Supplemental Results for "Noncoding and Coding Mechanisms of Aging Heart Failure with Preserved Ejection Fraction with Thyroid Dysfunction"

**Supplemental Figure S1: Diastolic dysfunction in early HFpEF**

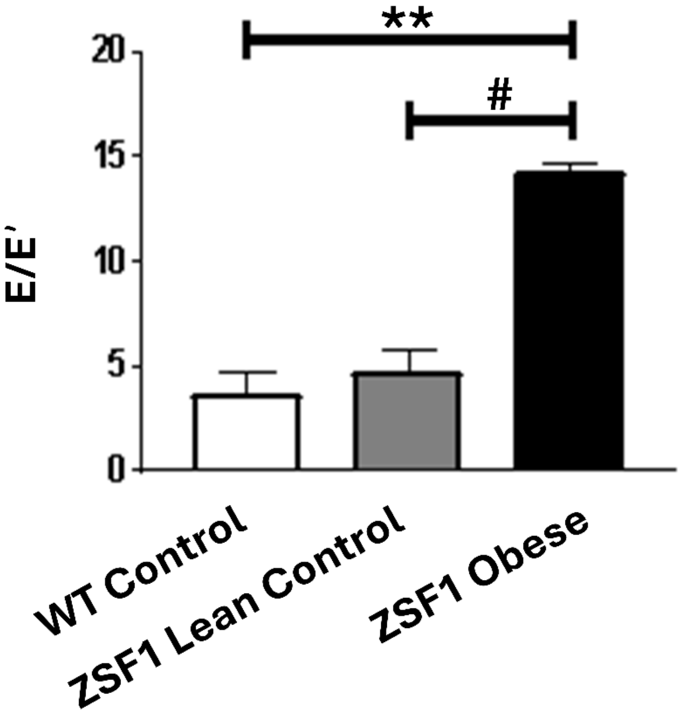


**Supplemental Figure S1: In early HFpEF rats, echocardiography showed significantly elevated E/E' indicating diastolic dysfunction.** All values are analyzed using one-way ANOVA and presented as means ± standard deviation; E/E': ratio of early mitral inflow velocity and mitral annular early diastolic velocity; WT: Wild Type; **$p<0.01$; [#]$p<0.05$.

**Supplemental Figure S2: Thyroid hormone levels in females (5 mo, 13 mo, 20 mo - WT control and ZSF1 lean control)**

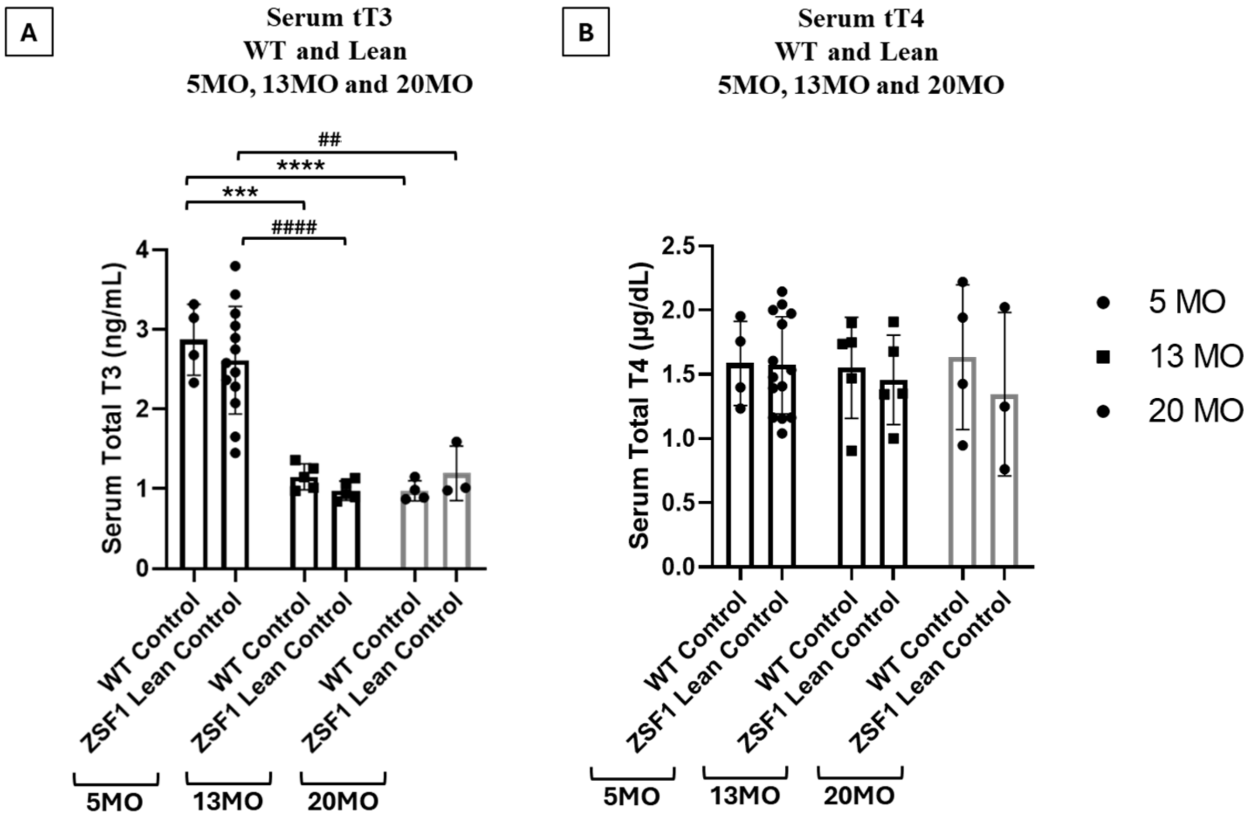


**Supplemental Figure S2: Serum thyroid hormone levels (Females):** Serum **(A)** total T3 and **(B**) total T4 levels in 5 mo, 13 mo, and 20 mo female WT control and ZSF1 lean control groups. All values are analyzed using Two-way ANOVA and presented as means ± standard deviation; WT: Wild Type; T4: Thyroxine; T3: Triiodothyronine; ***p<0.001, ****p<0.0001 vs 5 mo WT control; $^{\#\#}$p<0.01, $^{\#\#\#\#}$p<0.0001 vs 5 mo ZSF1 lean control.

**Supplemental Figure S3: Morphometrics in males (normalized by tibial length - 5 mo and 13 mo - WT control, ZSF1 lean control and ZSF1 obese HFpEF)**

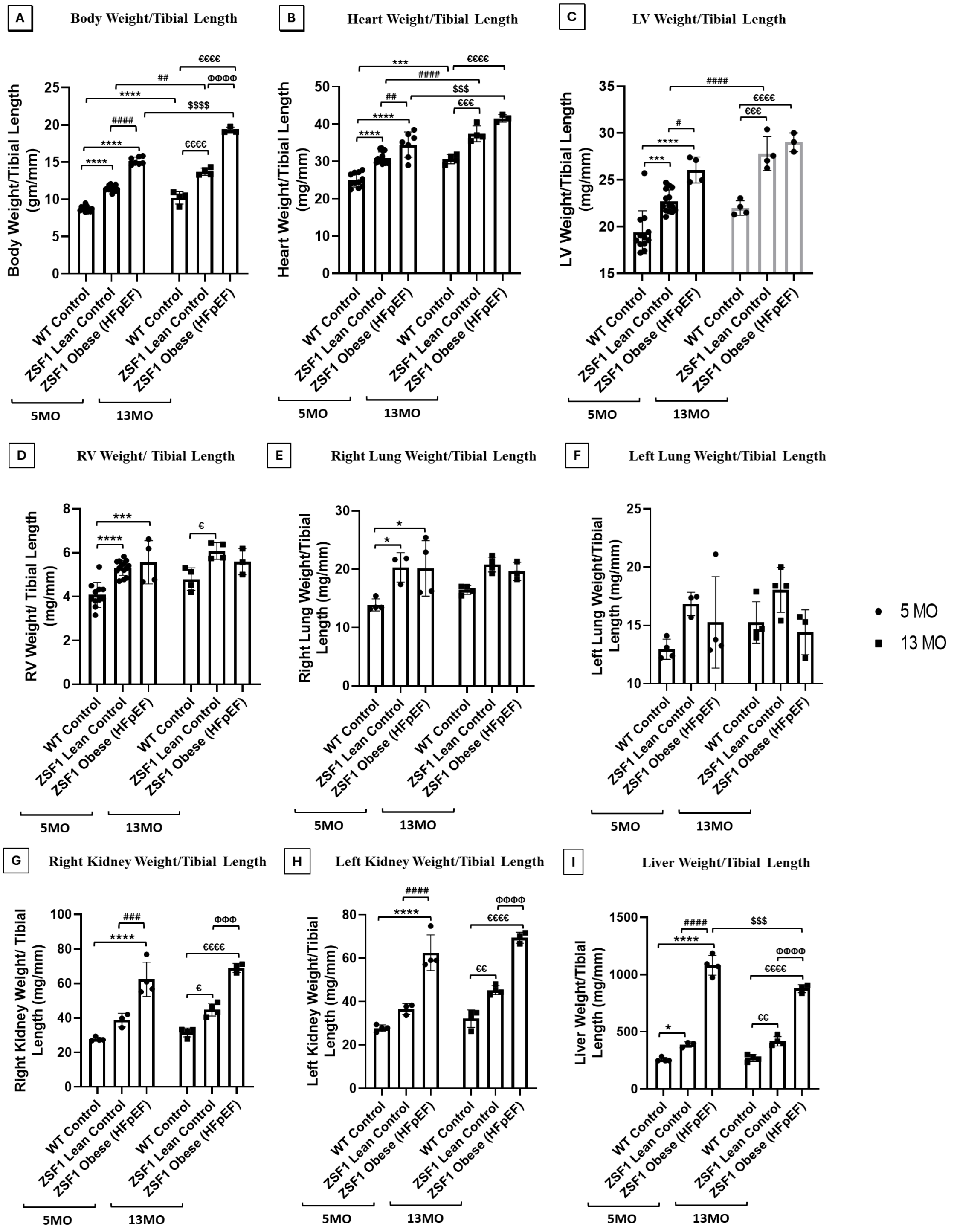

**Supplemental Figure S3: Morphometric analyses (Males)** of 5 mo and 13 mo WT control, ZSF1 lean control, and ZSF1 obese (HFpEF) rats **(A)** Body weight/Tibial Length; **(B)** Heart weight/Tibial Length; **(C)** LV weight/Tibial Length; **(D)** RV weight/Tibial Length; **(E)** Right lung weight/Tibial Length; **(F)** Left lung weight/Tibial Length; **(G)** Right kidney weight/Tibial Length; **(H)** Left kidney weight/Tibial Length and **(I)** Liver weight/Tibial Length. All values are analyzed using Two-way ANOVA and presented as means ± standard deviation; TH: thyroid hormone; WT: Wild-type, LV: Left ventricle, RV: Right ventricle weight, *$p<0.05$, ***$p<0.001$ ****$p<0.0001$ vs 5 mo WT control; $^{\#}p<0.05$, $^{\#\#}p<0.01$, $^{\#\#\#}p<0.001$, $^{\#\#\#\#}p<0.0001$ vs 5 mo ZSF1 lean control; $^{\$\$\$}p<0.001$, $^{\$\$\$\$}p<0.0001$ vs 5 mo ZSF1 obese (HFpEF); $^{\varepsilon}p<0.05$, $^{\varepsilon\varepsilon}p<0.01$, $^{\varepsilon\varepsilon\varepsilon}p<0.001$, $^{\varepsilon\varepsilon\varepsilon\varepsilon}p<0.0001$ vs 13 mo WT control; $^{\Phi\Phi\Phi}p<0.001$, $^{\Phi\Phi\Phi\Phi}p<0.0001$ vs 13 mo ZSF1 lean control.

**Supplemental Figure S4: Morphometrics in males (normalized by tibial length - 5 mo, 13 mo, 20 mo - WT control and ZSF1 lean control).**

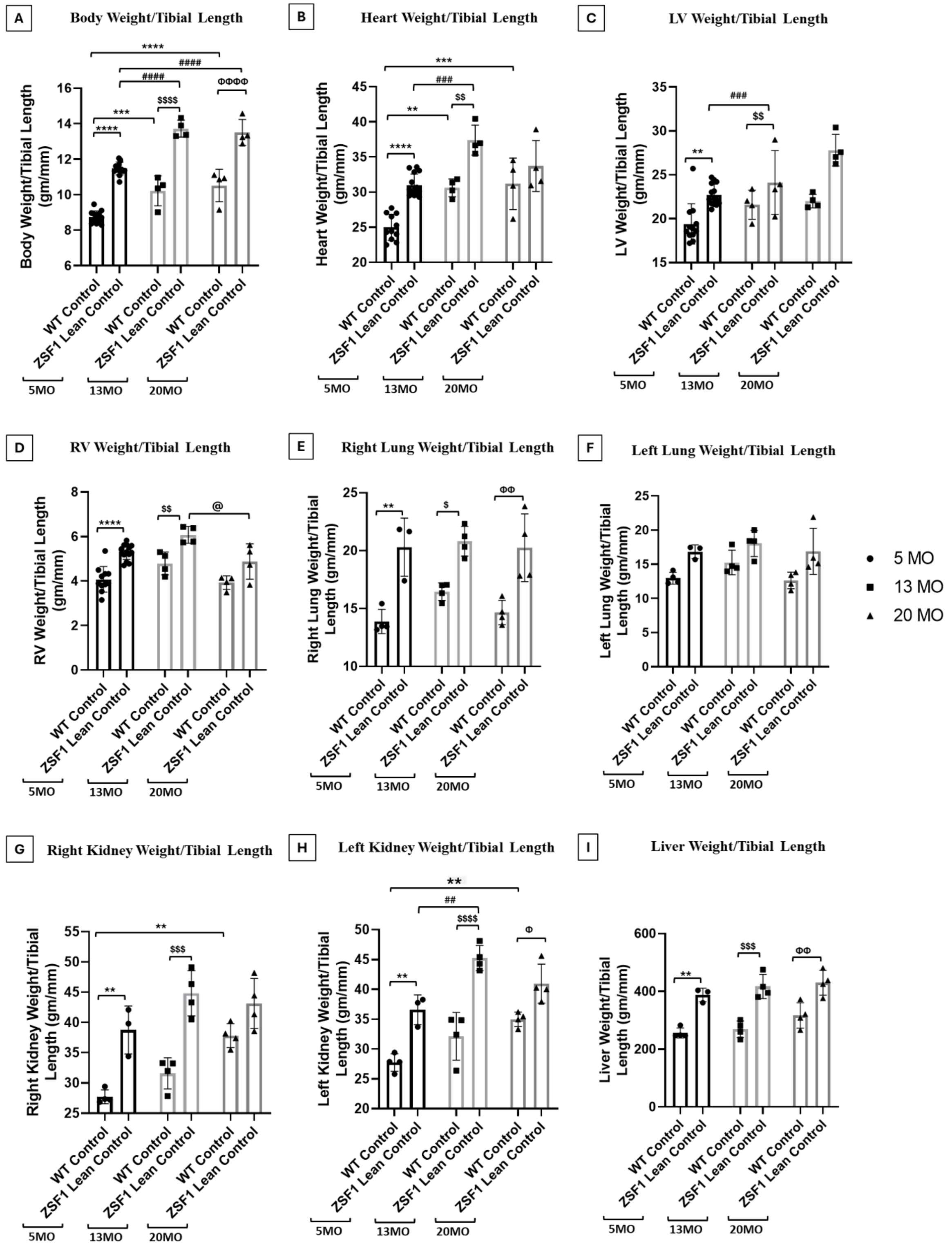

**Supplemental Figure S4: Morphometric analyses - Males** of 5 mo, 13 mo and 20 mo WT control and ZSF1 lean control rats (Male) **(A)** Body weight/Tibial length; **(B)** Heart weight/Tibial length; **(C)** LV weight/Tibial length; **(D)** RV weight/Tibial length; **(E)** Right lung weight/Tibial length; **(F)** Left lung weight/Tibial length; **(G)** Right kidney weight/Tibial length; **(H)** Left kidney weight/Tibial length and **(I)** Liver weight/Tibial length. All values are analyzed using Two-way ANOVA and presented as means ± standard deviation; WT: Wild-type, LV: Left ventricle, RV: Right ventricle weight; **p<0.01, ***p<0.001, ****p<0.0001 vs 5 mo WT control; $^{\#\#}$p<0.01, $^{\#\#\#\#}$p<0.0001 vs 5 mo ZSF1 lean control; $^{\$}$p<0.05, $^{\$\$}$p<0.01, $^{\$\$\$}$p<0.001, $^{\$\$\$\$}$p<0.0001 vs 13 mo WT control; $^{@}$p<0.05 vs 13 mo ZSF1 lean control; $^{\Phi}$p<0.05, $^{\Phi\Phi}$p<0.01, $^{\Phi\Phi\Phi\Phi}$p<0.0001 vs 20 mo WT control.

**Supplemental Figure S5: Morphometrics in females (5 mo, 13 mo, 20 mo - WT and ZSF1 lean control)**

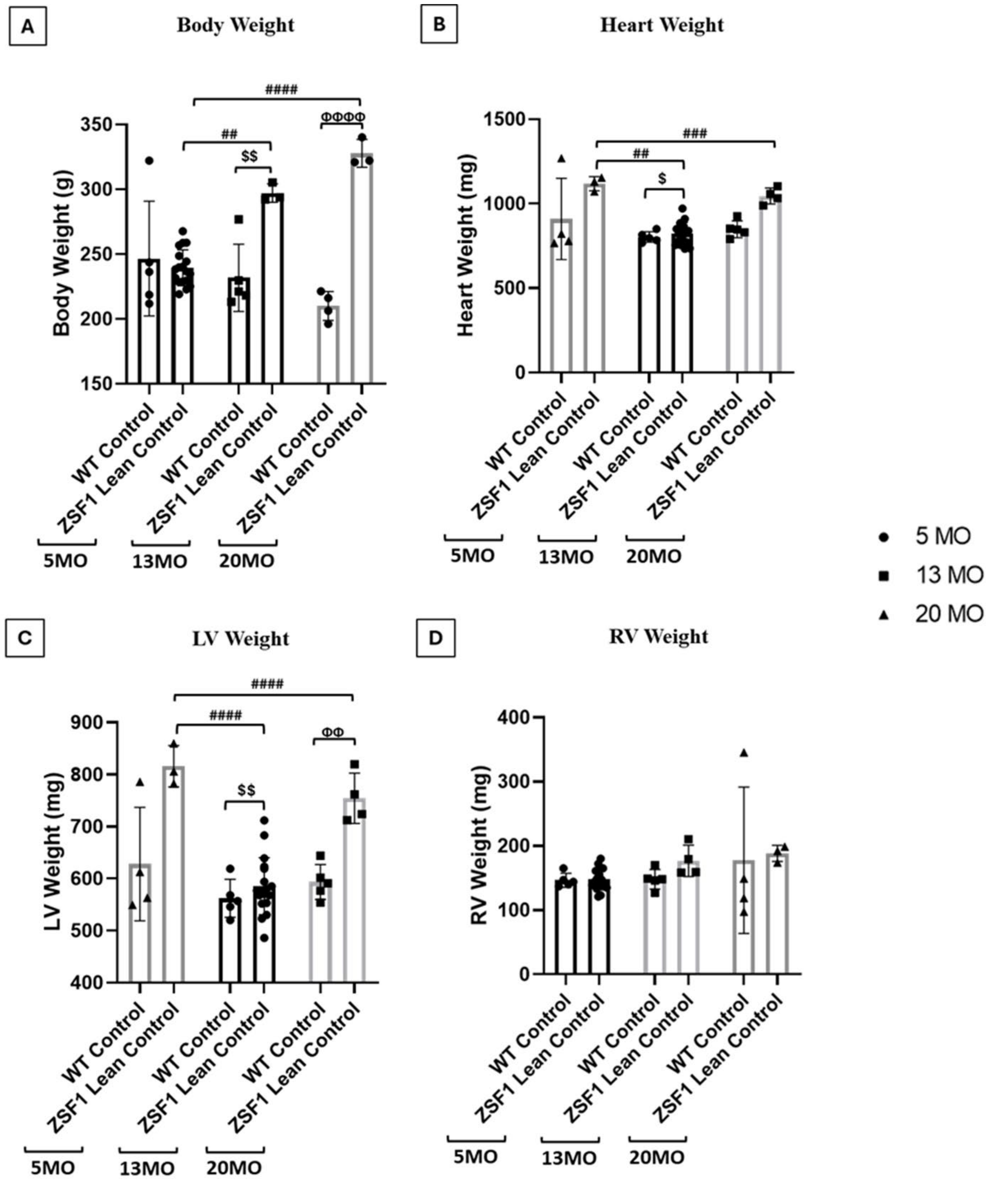

**Supplemental Figure S5: Morphometric analyses - Females** of 5 mo, 13 mo, and 20 mo WT control and ZSF1 lean control rats **(A)** Body weight **(B)** Heart weight **(C)** LV weight and **(D)** RV weight. - All values are analyzed using Two-way ANOVA and presented as means ± standard deviation; WT: Wild-type, LV: Left ventricle, RV: Right ventricle weight; $^{\#\#}p<0.01$, $^{\#\#\#}p<0.001$, $^{\#\#\#\#}p<0.0001$ vs 5 mo ZSF1 lean control; $^{\$}p<0.05$, $^{\$\$}p<0.01$ vs 5 mo ZSF1 obese (HFpEF); $^{\theta\theta}p<0.01$, $^{\theta\theta\theta\theta}p<0.0001$ vs 13 mo ZSF1 lean control.

**Supplemental Figure S6: Morphometrics in females (normalized by tibial length; 5 mo, 13 mo, 20 mo - WT and ZSF1 lean control)**

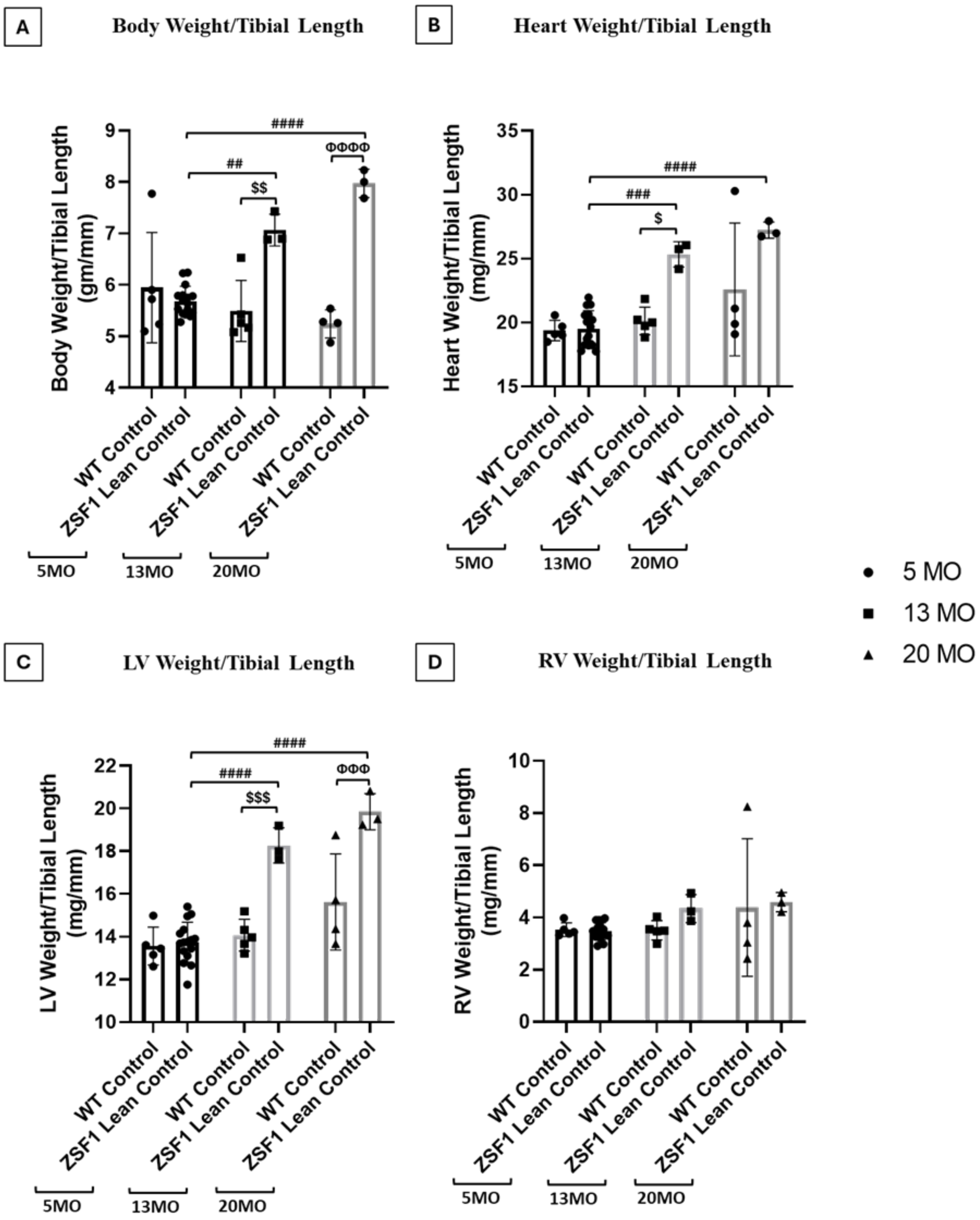

**Supplemental Figure S6: Morphometric analyses (Female)** of 5 mo, 13 mo, and 20 mo WT control and ZSF1 lean control rats **(A)** Body weight/Tibial Length **(B)** Heart weight/Tibial Length **(C)** LV weight/Tibial Length and **(D)** RV weight/Tibial Length. All values are analyzed using Two-way ANOVA and presented as means ± standard deviation; WT: Wild-type, LV: Left ventricle, RV: Right ventricle weight; $^{\#\#}p<0.01$, $^{\#\#\#}p<0.001$, $^{\#\#\#\#}p<0.0001$ vs 5 mo ZSF1 lean control; $^{\$}p<0.05$, $^{\$\$}p<0.01$, $^{\$\$\$}p<0.001$ vs 5 mo ZSF1 obese (HFpEF); $^{\Phi\Phi\Phi}p<0.001$, $^{\Phi\Phi\Phi\Phi}p<0.0001$ vs 20 mo WT control.
